# *In vitro* and *in vivo* analysis of microvesicle-mediated metastasis using a bright, red-shifted bioluminescent reporter protein of extracellular vesicles

**DOI:** 10.1101/2021.07.23.453512

**Authors:** Ahmed A. Zarea, Gloria I. Perez, David Broadbent, Benedikt Dolgikh, Matthew P. Bernard, Alicia Withrow, Amelia McGill, Victoria Toomajian, Lukose K. Thampy, Jack Harkema, Joel R. Walker, Thormas A. Kirkland, Michael H. Bachmann, Jens Schmidt, Masamitsu Kanada

## Abstract

Cancer cells produce heterogeneous extracellular vesicles (EVs) as mediators of intercellular communication. Our study focused on a novel method to image EV subtypes and their biodistribution *in vivo*. Regardless of injection routes, we established that reporter EVs isolated from murine mammary carcinoma cells expressing PalmReNL, which utilizes bioluminescence resonance energy transfer (BRET), localized to the lungs. This new EV reporter allowed highly sensitive EV tracking *in vitro* and *in vivo* and enabled us to begin studies to understand the commonalities and functional differences of the EV subtypes. We demonstrated the early appearance of metastatic foci in the lungs of mammary tumor-bearing mice following multiple injections of the microvesicle (MV)-enriched fraction derived from mammary carcinoma cells. In addition, the results we present here show that tumor cell-derived MVs act on distant tissues through upregulating LC3 expression within the lung.

## Introduction

Extracellular vesicles (EVs) are spherical lipid bilayered structures naturally shed by cells and have been implicated in the pathogenesis of cancer and numerous other diseases^1,2^. Understanding their biodistribution and ultimate targets is key to elucidating their roles in health and disease^3,4^. EV subtypes include exosomes and microvesicles (MVs), which are distinguished based on their size and biogenesis^1,2^. Exosomes range from ∼30-120 nm in diameter and are produced by inward budding of the late endosomal membrane, known as multivesicular bodies (MVBs). MVs are 50-1000 nm in diameter and produced by simple outward budding of the plasma membrane. Due to their nano-size and biophysical properties, both types of EVs have the potential to cross biological barriers and gain access into host cells beyond these barriers^5–7^. In this manner, released EVs act as mediators of intercellular communication in the body^8,9^. For example, numerous studies have demonstrated that cancer cells can appropriate this communication pathway by transferring active biomolecules to adjacent and distant cancer cells, promoting their growth and survival^2,10^. For this reason, EV-mediated signaling may hold promising cancer treatment strategies and be an effective platform for drug delivery. However, systemic administration of nano-sized EVs may reach and accumulate in other sites beyond the tissues of therapeutic interest^11–13^. Therefore, analysis of the biodistribution following EV administration is a prerequisite for the development of EV-based therapeutics.

Characterization of EV biodistribution, however, is restricted by the biological tools available, which are not sensitive enough to localize and track small EVs *in vivo*. In this study, we develop a novel EV reporter system that enables highly sensitive EV tracking. Non-invasive *in vivo* bioluminescence imaging combined with molecular and cellular analyses offers unique potential to facilitate the preclinical evaluation of biological therapies in animal models. Additionally, most *in vivo* studies of EV-mediated signaling have only used immune-compromised mice^3,14^ and require further assessment in the presence of an intact immune system. In our current studies, combining *in vitro* and *in vivo* approaches with immunocompetent mice, we expect to uncover some of the biological characteristics of EVs, such as trafficking, cellular uptake and release.

## Results

### Overexpression of PalmReNL labeled both exosomes and MVs

We developed PalmReNL, a novel EV imaging probe, by genetically fusing a palmitoylation signal peptide^15^ to one of the brightest red-shifted bioluminescence resonance energy transfer (BRET) reporters, Red-eNanoLantern (ReNL)^16^ (Fig. 1a). We observed that PalmReNL supplied with its substrate furimazine (Fz) produced red-shifted luminescence similar to that of ReNL without palmitoylation, indicating that membrane-anchoring of ReNL does not affect its BRET efficiency (Fig. 1b). This EV reporter has an advantage for *in vivo* tracking since bioluminescence imaging produces negligible background signals. Also, photons with spectral wavelengths longer than 600 nm can efficiently penetrate mammalian tissues with less light attenuation than observed with shorter wavelength light. Moreover, PalmReNL can be used as a fluorescent EV reporter by exciting tdTomato for standard flow cytometry and microscope settings.

**Fig. 1.**
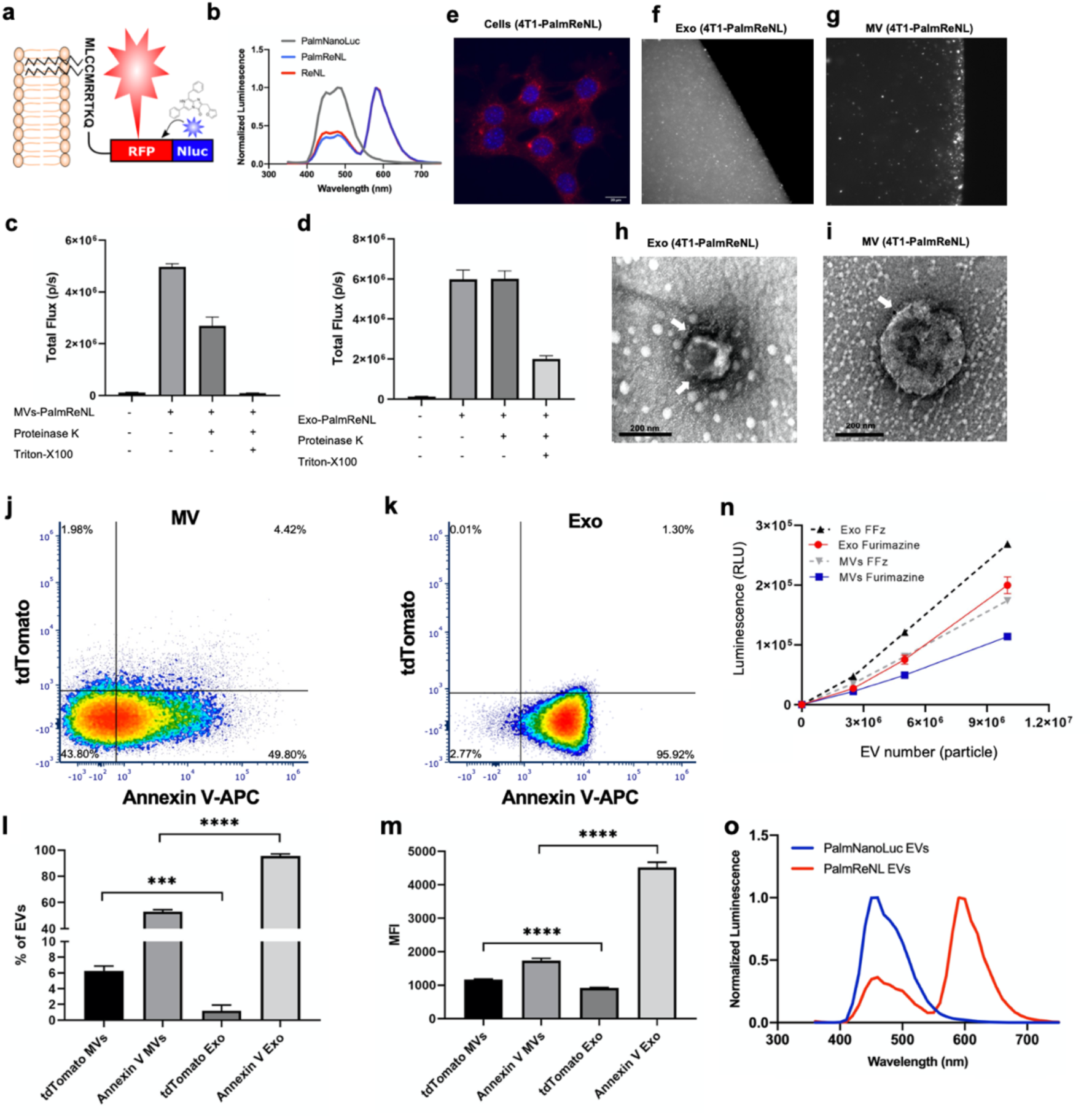
PalmReNL-based labeling of exosomes and MVs derived from 4T1 cells. **a** Schematic diagram of EV membrane labeling with PalmReNL BRET probe. **b** Emission spectra of murine mammary carcinoma 4T1 cells transfected with ReNL, PalmReNL, or PalmNanoLuc. **c, d** Proteinase-K protection assay for PalmReNL-MVs and exosomes. **e** 4T1 cells constitutively expressing PalmReNL. Punctate signals of RFP (red) were merged with nuclei stained with Hoechst 33342. Scale bar, 20 µm. **f, g** A droplet of buffer containing isolated PalmReNL-exosomes and -MVs. **h, i** Transmission electron microscopy of 4T1 cell-derived PalmReNL-exosomes or -MVs, immunogold labeled for RFP. Arrows point towards positive RFP signal (dark spots). Scale bars, 200 nm. **j, k** A representative of three independent experiments of Annexin V staining of individual PalmReNL-MVs and -exosomes analyzed by flow cytometry. The tdTomato fluorescence signal represents PalmReNL-EVs. FACS plots were gated for tdTomato^+^ and Annexin V^+^ EVs among the CTV-stained EVs. **l, m** The percentage and median fluorescence intensity (MFI) of positively labeled CTV-exosomes and - MVs. Error bars, SD (n = 3), ***, P < 0.001; ****, P < 0.0001. **n** Bioluminescence analysis of PalmReNL-exosomes and MVs using furimazine and fluorofurimazine (FFz). Error bars, SD (n = 5). **o** Emission spectra of the isolated PalmNanoLuc- and PalmReNL-EVs.

As reported previously^15^, the results of the experiments to determine the membrane orientation of PalmReNL in EVs, revealed that PalmReNL can label the inner membrane leaflet of both exosomes and MVs. However, there were differences in the outer membrane labeling by PalmReNL between exosome- and MV-enriched fractions. As demonstrated by both the dot blot (Supplementary Fig. 1f-i) and the proteinase-K protection assays (Fig. 1c, d), in exosomes the reporter was protected from Proteinase-K, hence localized primarily to the inner membrane. In contrast, in MVs PalmReNL was sensitive to Proteinase-K without detergent treatment, indicating that it localized to both the inner and the outer membranes. This dual-labeling of exosomes and MVs represents an advantage over other widely used CD63-based EV reporters that only label one specific EV subtype, exosomes^17^.

We next assessed the labeling efficiency in isolated exosome- and MV-enriched fractions from 4T1 cells stably expressing PalmReNL. Firstly, exosomes and MVs were isolated from the conditioned medium as we previously reported^18,19^ and characterized by nanoparticle tracking analysis (NTA). The concentration of MVs expressing PalmReNL was 7.7 × 10^10^ particles/mL and the mean diameter was 119 nm (Supplementary Fig. 1b), while the concentration of exosomes expressing PalmReNL was 1.9 × 10^10^ particles/mL and the mean diameter was 105 nm (Supplementary Fig. 1a). The concentrations and mean diameters of exosome- and MV-enriched fractions derived from control 4T1 cells were 1.9 × 10^10^ particles/mL and 102 nm, 2.7 × 10^10^ particles/mL and 117 nm, respectively (Supplementary Fig. 1c, d). Therefore, the genetic addition of the reporter did not inhibit the release of EVs nor influence their sizes. Determination of the Zeta potential revealed that the PalmReNL slightly shifted the surface charge of exosomes, but not MVs (Supplementary Fig. 1e). Consistent with the NTA data, the transmission electron microscopy (TEM) analysis of the exosome- and MV-enriched fractions revealed a heterogeneous mixture of predominantly intact vesicles with artefactual cup-shaped morphology^20^ having diameters ranging from 50 to 200 nm, and expressing the tdTomato as ReNL is a fusion protein of NanoLuc and tdTomato^16^ (Fig. 1h, i). There was no significant morphological change in the EV fractions expressing PalmReNL.

Western blot (WB) analysis of exosome marker proteins in immunoblots of whole-cell lysates and EVs derived from 4T1 cells expressing PalmReNL (Supplementary Fig. 1k) demonstrated that exosome fractions preferentially express CD63, TSG101, and Alix, whereas cell lysates and MV-enriched fractions preferentially express Flotillin-1 as we previously reported^19^. Importantly, WB analysis of EVs collected from the parental 4T1 cell line demonstrated that the reporter protein (PalmReNL) does not interfere with the expression of any of the EVs marker proteins tested (Supplementary Fig. 1j). All fractions of EVs containing PalmReNL (cell lysates, MVs, and exosomes) expressed tdTomato. The labeling efficiency was also confirmed by fluorescence microscopy, demonstrating that the total fluorescence intensities were higher in exosomes, but punctate signal intensity was higher for individual MVs (Fig. 1e-g). To further assess the efficiency of EV labeling with PalmReNL, we analyzed individual PalmReNL-exosomes and -MVs by flow cytometry (Fig. 1j, k). Firstly, all the isolated PalmReNL-EVs were stained with CellTrace Violet (CTV) as an alternative to CFSE, an amine-reactive dye previously used for nanoFACS^21^. The PalmReNL signal was detected on the tdTomato channel, and the percentage of positive labeling was 1.2% for exosomes and 6.3% for MVs stained with CTV (Fig. 1l). The median fluorescence intensity (MFI) of PalmReNL in individual MVs was 1.27-fold higher than that of exosomes (Fig. 1m). As we previously reported, phosphatidylserine (PS) externalization in exosomes and MVs was examined using Annexin V staining^18^. The fluorescence signals in individual exosomes were significantly higher than the signals in individual MVs, where 95.4% and 52.9% of PS externalization was detected in exosomes and MVs stained with CTV, respectively (Fig. 1l). The MFI of Annexin V in individual MVs was 2.6-fold lower than that of exosomes among the CTV-stained EVs (Fig. 1m).

Furthermore, we found that the release of EVs from 4T1 cells-expressing PalmReNL *in vitro* occurs every 24 h and peaks at 72 h. The bioluminescence signals in both exosomes- and MVs-PalmReNL correlated with the number of particles present, and increased steadily over the 72 h period (Supplementary Fig. 2b, c). The reporter 4T1 cells appear to release ∼20-fold more MVs than exosomes as assessed by NTA. The bioluminescence signals were also correlated with the number of EVs purified from different EV fractions after density gradient ultracentrifugation (Supplementary Fig. 2d). Measuring the bioluminescence signals in equal numbers of PalmReNL-exosomes and - MVs ranging from 2.5 × 10^6^ to 1.0 × 10^7^ using 25 µM Fz demonstrated that the bioluminescence signals in the exosomes (1.9 × 10^5^ ± 1.4 × 10^4^ RLU; *p*<0.0001) were 1.7-fold higher than those of the MVs (1.1 × 10^5^ ± 1.8 × 10^3^ RLU). A novel Fz analogue, named fluorofurimazine (FFz), with increased aqueous solubility was recently developed^22^. We found FFz was 1.4- and 1.5-fold more sensitive (exosomes 2.6 × 10^5^ ± 3.9 × 10^3^ RLU, *p*=0.0012; MVs 1.7 × 10^5^ ± 3 × 10^3^ RLU, *p*<0.0001) than Fz (Fig. 1n) with PalmReNL-exosomes and -MVs, respectively. The protein concentrations of 5.6 × 10^7^ EV particles were 45.4 µg/mL for exosomes and 23.9 µg/mL for MVs. Furthermore, the bioluminescence spectra of PalmReNL-exosomes and -MVs were measured with similar emission spectra, peaking at 585 nm (Fig. 1o). Taken together, these results suggest that exosomes incorporate PalmReNL more efficiently compared to MVs. However, fluorescence signals in individual exosomes are below the detection limit and only individual MVs carry fluorescently detectable numbers of PalmReNL molecules due to the larger surface areas relative to exosomes and/or symmetrical labeling of MV membranes.

### Rates of endocytosis of tumor cell-derived exosomes and MVs were similar between various recipient cell types

Uptake of PalmReNL-exosomes and MVs by macrophages (Fig. 2a-c), 4T1 cells (Fig. 2d-f), lung fibroblasts (Fig. 2g-i), or adipose-derived mesenchymal stromal cells (AMSCs) were assessed (Fig. 2j-l). Phase contrast and fluorescence microscopy demonstrated that EV-uptake was time-dependent. In our experimental conditions the 24 h time point showed the highest fluorescence. Of note, the fluorescence signals of our reporter EVs decreases over time, which is an advantage compared to the widely used lipophilic fluorescent dyes such as PKH, DiR, and DiI which can persist in the recipient cells. These dyes previously showed inaccurate spatiotemporal information on the reporter EVs due to its inherent stability^15,18^.

**Fig. 2.**
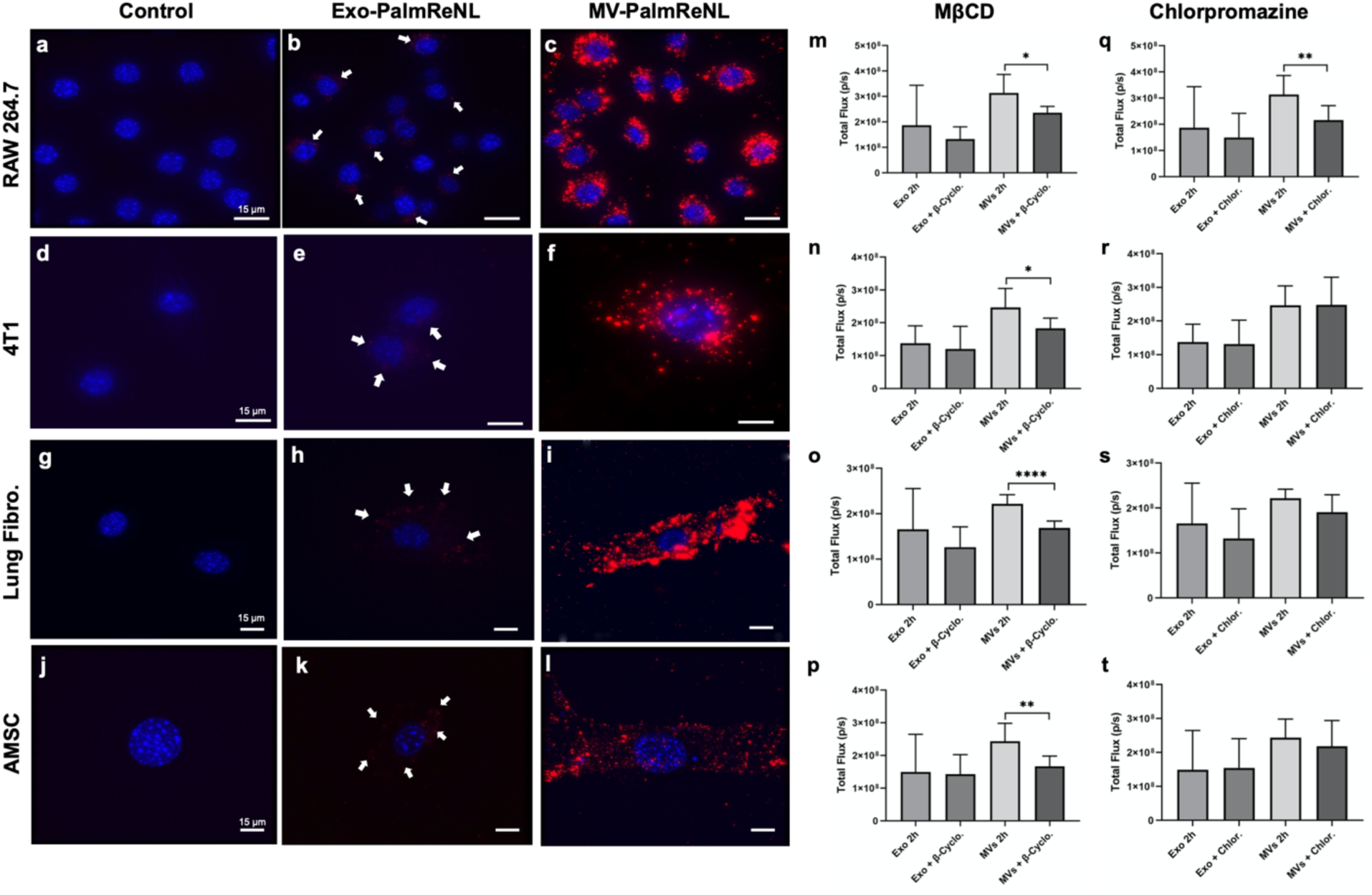
Similar rates of endocytosis of 4T1 cell-derived exosomes- and MVs- PalmReNL between various recipient cell types *in vitro*. **a-c** Macrophages (RAW 264.7). **d-f** 4T1 cells. **g-i** primary mouse lung fibroblasts. **j-l** Adipose-derived mesenchymal stromal cells (AMSCs). Punctate signals of RFP (red) were merged with nuclei stained with Hoechst 33342 (blue). Scale bar, 15 µm. Arrows indicate weak RFP signals in PalmReNL-exosomes. **m-t,** The recipient cells were treated with methyl-β-cyclodextrin (MβCD; 10 mM) (m-p) or Chlorpromazine (10 ug/mL) (q-t). **m, q** Macrophage RAW 264.7 cells. **n, r** 4T1 cells. **o, s** Mouse primary lung fibroblasts. **p, t** Mouse AMSCs. Error bars, SD (n = 8), *, P < 0.05; **, P < 0.01; ****, P < 0.0001.

The bioluminescence signal did not reveal any significant differences between the cell types in the uptake of the reporter, except at 2 h between RAW 264.7 cells and lung fibroblasts where the bioluminescence signal of PalmReNL-MVs was significantly higher (3.1 × 10^8^ ± 2.5 × 10^7^ and 2.2 × 10^8^ ± 7.1 × 10^6^ p/s, respectively; *p*=0.004). Both cell types had a significantly lower (*p*<0.05) bioluminescence signal of PalmReNL-MVs at 24 h (Supplementary Fig. 3a-d). By contrast, the uptake of PalmReNL-exosomes appeared to remain constant during the 24 h period in all cell types tested. Next, we assessed the mechanism of cellular uptake of exosomes and MVs. Inhibition of caveolin-dependent endocytosis, which was inhibited by methyl-β-cyclodextrin (MβCD; a compound that sequesters the cholesterol in the cell membrane^23^), significantly decreased the MV-uptake in all cell types studied (Fig. 2m-p). Interestingly, MV uptake in Raw 264.7 cells appears to occur by both clathrin-dependent endocytosis inhibited by chlorpromazine^24^ and caveolin-dependent endocytosis (Fig. 2m, q). On the other hand, the uptake of exosomes in all the cell types tested was independent of both caveolin and clathrin (Fig. 2m-t).

Although the bioluminescence signal for PalmReNL-exosomes was higher than that of MVs-PalmReNL (see Fig. 1n), detecting exosome uptake by fluorescence microscopy was challenging. We hypothesized that our inability to detect the PalmReNL-exosomes reflects those reporter exosomes cannot retain the fluorescence signal after being taken up by the cells. Rapid diffusion of the PalmReNL into other cellular membrane compartments might result in a fast loss of signals. To prove this, we compared the *in vitro* uptake by RAW 264.7 macrophages or 4T1 cells of exosomes derived from 4T1 cells stably expressing the exosome marker CD63 fused with mScarlet^25^ or PalmReNL. CD63-mScarlet-exosomes showed punctate fluorescence signals in both recipient 4T1 and RAW 264.7 cells (Fig. 3c, f), demonstrating their cellular uptake and signal retention after being taken up by cells. On the other hand, PalmReNL-exosomes did not retain the signals after being taken up by cells and therefore precluded their visualization with the current sensitivity and resolution of our microscope (Fig. 3b, e). However, exosomes carrying PalmReNL retained bioluminescence in the recipient cells (see Supplementary Fig. 3). These results indicate that PalmReNL carried by exosomes might be rapidly transfered from early endosomes into other membrane compartments for either degradation or recycling, whereas transferred CD63-mScarlet may be retained in endosomal membranes in recipient cells as recently reported^17^.

**Fig. 3.**
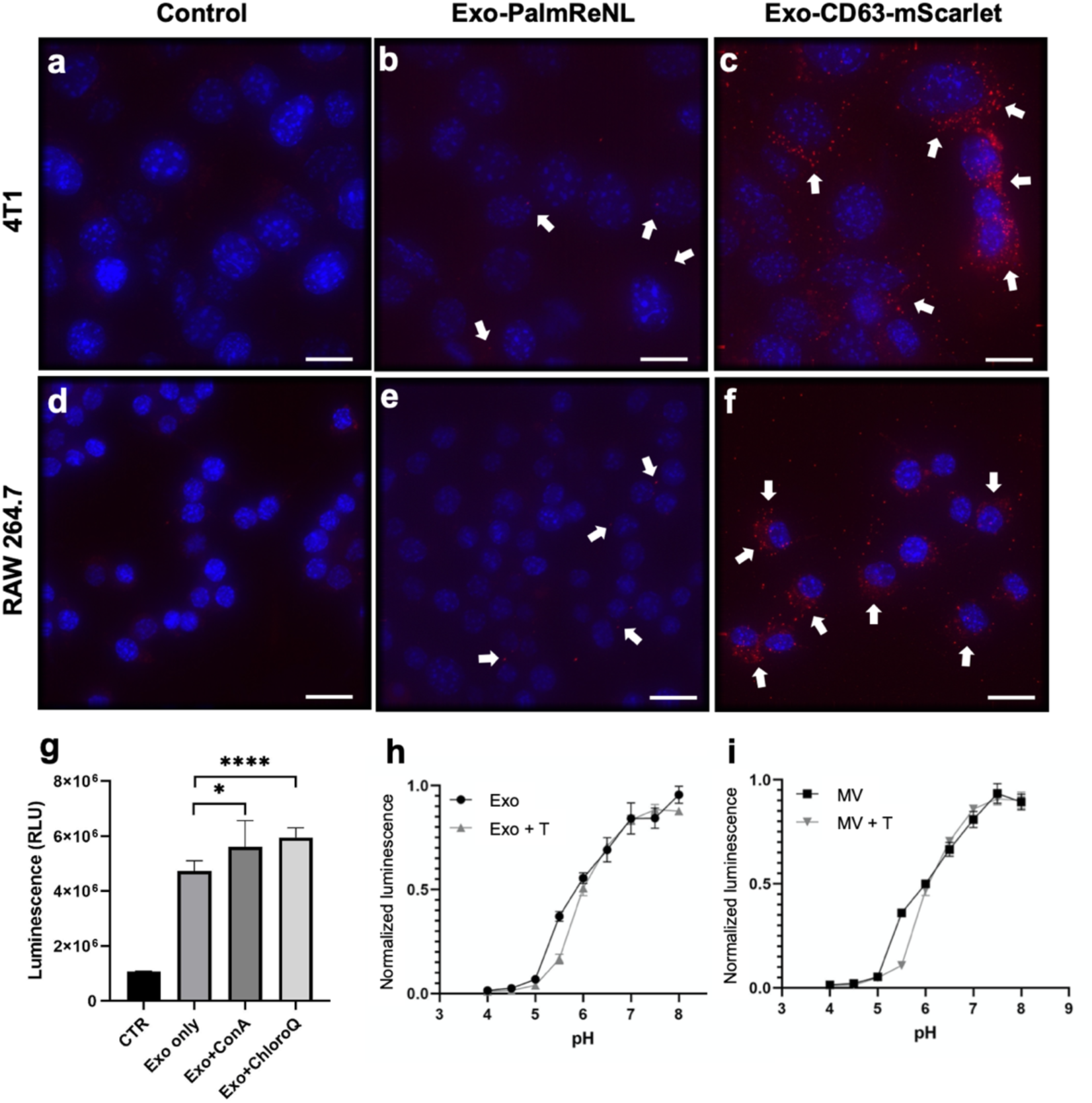
Rapid processing of PalmReNL carried by exosomes via the endosomal-lysosomal pathway. Fluorescence microscopy images of 4T1 and RAW 264.7 cells treated for 24 h with PalmReNL- or CD63-mScarlet-exosomes. **a, d** Control 4T1 and RAW 264.7 cells. **b, e** 4T1 and RAW 264.7 cells treated with exosomes-PalmReNL. **c, f** 4T1 and RAW 264.7 cells treated with CD63-mScarlet-exosomes. Punctate fluorescence signals were merged with nuclei stained with Hoechst 33342. Scale bar, 15 µm. White arrows, PalmReNL- or CD63-mScarlet-exosomes. **g** PalmReNL-exosomes taken up by 4T1 cells in the presence of concanamycin-A or Chloroquine showed higher bioluminescence signals compared to control. Error bars, SD (n = 8), *, P < 0.03, ****, P < 0.0001. **h, i** Conventional pH titration curves of the normalized bioluminescence signals of exosomes- and MVs-PalmReNL. Error bars, SD (n = 5).

To further assess the effect of acidic cellular compartments on PalmReNL-exosomes, we used Palm-fused Gamillus (acid-tolerant monomeric GFP^26^) for EV labeling and compared the uptake of exosomes by fluorescence microscopy. At 24 h the exosomes-PalmGamillus signal was easily detected by fluorescence microscopy compared to the PalmReNL-exosomes signal which was barely detected (Supplementary Fig. 4b, c). Moreover, treatment of cells with inhibitors of endosomal acidification, either Concanamycin A^27^ or Chloroquine^28^ significantly increased the bioluminescence signal of PalmReNL-exosomes (Fig. 3g). Since the loss of NanoLuc activity with endosomal translocaiton was previously reported^29^, we analyzed the pH sensitivity of EVs-PalmReNL. The bioluminescence signals of both PalmReNL-exosomes and -MVs steadily decreased at pH below 6.0 either with or without detergent treatment (Fig. 3h, i), indicating significant signal loss of PalmReNL-based EV reporters in acidic cellular compartments.

### Proliferation of various cell types by treatment with EVs *in vitro*

To evaluate the physiological significance of cellular EV-uptake we analyzed the proliferation curves of various cell types (Raw 264.7, 4T1, lung fibroblasts, and AMSCs) when cultured for a period of 48 h with or without EVs (2.5 × 10^9^) derived from 4T1 cells expressing PalmReNL. Interestingly, only the exosome-enriched fraction increased the proliferation rate of every one of the cell types tested. By contrast, MVs increased the proliferation rate only in RAW 264.7 cells (Fig. 4b). Both exosomes and MVs activated macrophages (clearly evident by the change in morphology of cell spreading and formation of dendrite-like structures) after 48 h of treatment (compare Fig. 4d, e with Fig. 4c).

**Fig. 4.**
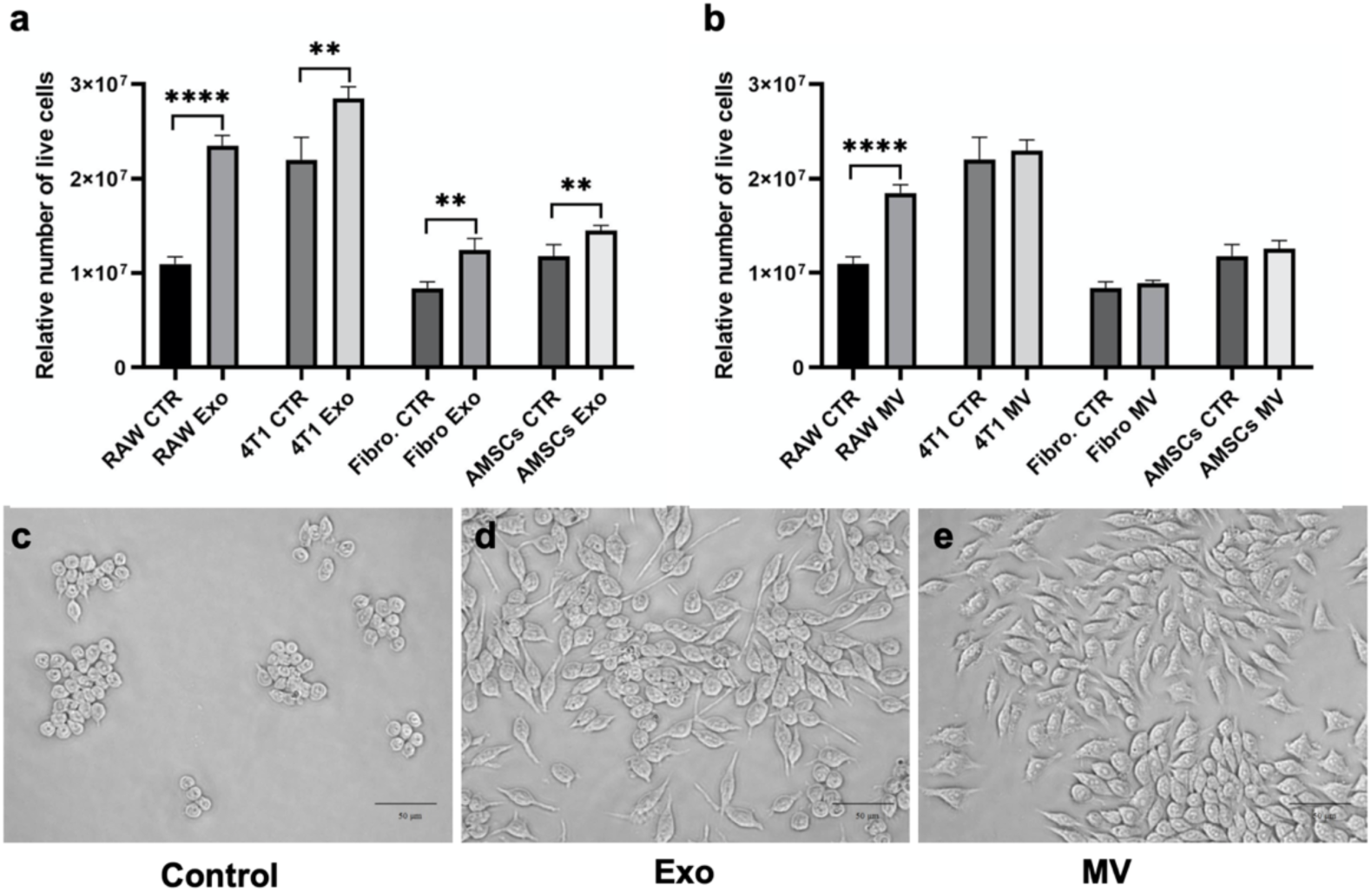
Proliferation of various cell types by treatment with 4T1 cell-derived EVs *in vitro.* **a, b** Proliferation curves of various cell types (RAW 264.7 cells, 4T1 cells, lung fibroblasts and AMSCs) when cultured for a period of 48 h without or with EVs-PalmReNL (2.5 × 10^9^). Error bars, SD (n = 4), **, P < 0.001; ****, P < 0.0001. **c, d, e** Both exosomes and MVs activated macrophages (clearly evident by the change in morphology) after 48 h of treatment. Scale bar, 50 µm.

### Exosomes and MVs derived from metastatic mammary carcinoma 4T1 cells showed similar biodistribution and preferentially accumulated in the lung *in vivo*

To determine the biodistribution of 4T1 cell-derived exosomes and MVs, 1.0 × 10^9^ exosomes or MVs carrying PalmReNL were administered intravenously (i.v.) in healthy female BALB/c mice. Both reporter exosomes and MVs distributed throughout the body within five min despite the difference of size and membrane composition between these EV classes. The reporter exosomes displayed significantly higher bioluminescence signals (1.8 × 10^6^ ± 8.8 × 10^5^ p/s; n=18; *p*=0.0016; Fig. 5b, c, f) following i.v. injections, compared to the reporter MVs i.v. injected (1 × 10^6^ ± 6.2 × 10^5^ p/s; n=15; Fig. 5d, e, f). The *ex vivo* signal, particularly in the lungs, was higher for the reporter exosomes (3.0 × 10^7^ ± 2.4 × 10^7^ p/s; n=3; *p*=0.02; Fig. 5g), when compared to signal in lungs from control mice (3.9 × 10^4^ ± 3.3 × 10^4^ p/s; n=5; Fig. 5g). However, there was no significant difference between the lung *ex vivo* signal for the reporter exosomes compared to that of the MVs (Fig. 5g). Moreover, there were no significant differences between the bioluminescence signals of the reporter exosomes and MVs injected i.p. (Fig. 5f).

**Fig. 5.**
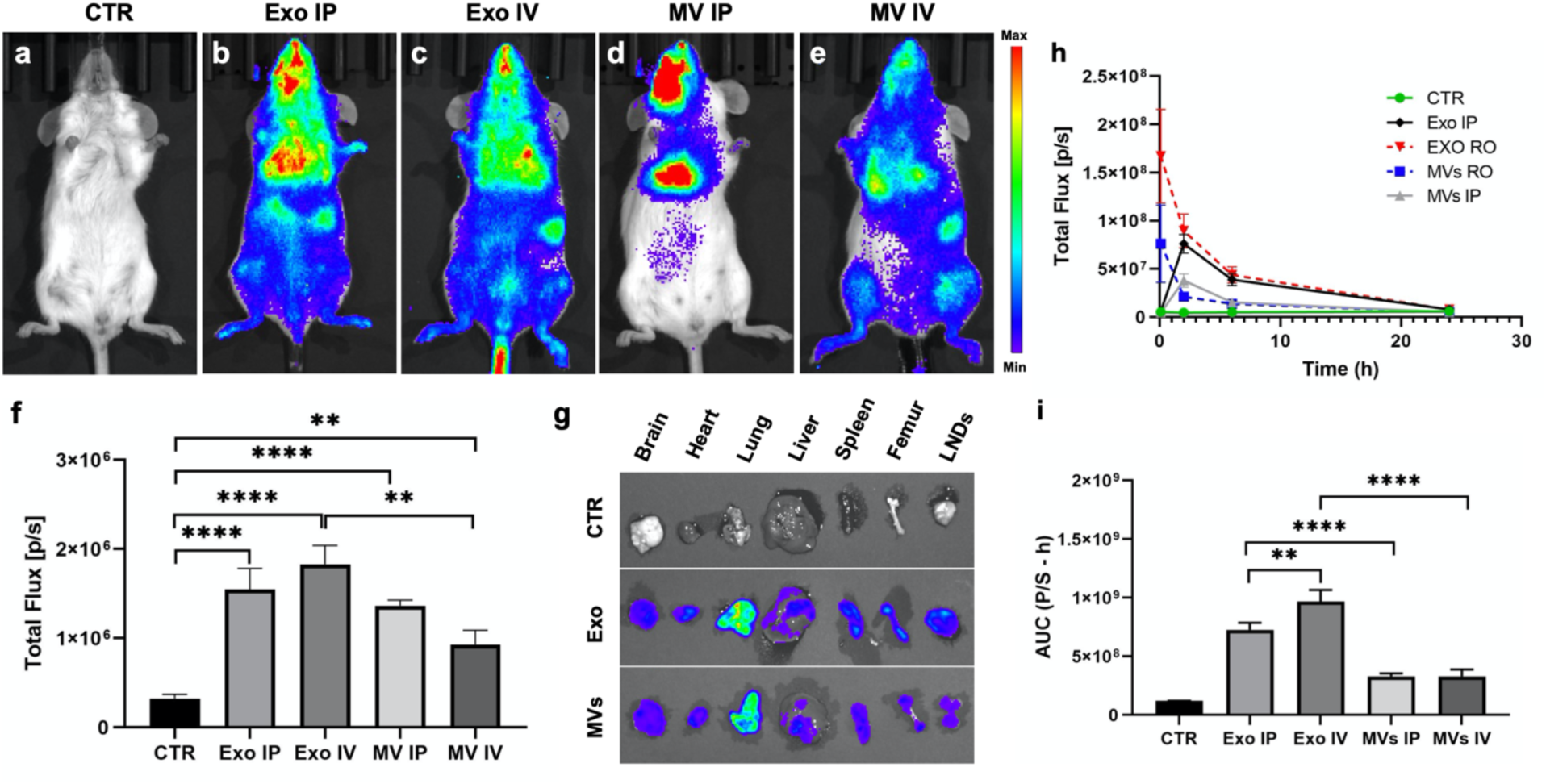
Similar biodistribution and organotropism of exosomes and MVs derived from metastatic mammary carcinoma 4T1 cells. **a-e** 4T1 cell-derived exosomes and MVs, 1.0 × 10^9^ PalmReNL-exosomes or -MVs were administered intravenously (i.v.) or intraperitonealy (i.p.) in healthy BALB/c mice. Furimazine was i.v. injected. **f** Analysis of bioluminescence signals in (a-e). There were no significant differences between the bioluminescence signals of PalmReNL-exosomes and -MVs injected i.p. **g** The *ex vivo* signal, particularly in the lungs, was higher for the reporter exosomes when compared to the signal in lungs from control mice. Error bars, SEM (CTR n =10; exosome i.p. n=3; exosome i.v. n=18; MV i.p. n=3; MV i.v. n=15), **, P < 0.01; ****, P < 0.0001. **h** The plasma samples were collected from the animals after EV isolation. The reporter exosomes displayed significantly higher bioluminescence signals compared to the reporter MVs in the plasma samples. The maximum bioluminescence signals were observed at 5 min in the i.v. injected reporter exosomes and MVs. In contrast, it took 2 h to reach the maximum levels of the bioluminescence signals in the i.p. injected reporter exosomes and MVs. Error bars, SD (n = 3). **i** AUC analysis of the bioluminescence signals in the plasma samples over 24 h post-EV injection. Error bars, SD (n = 15), **, P < 0.01; ****, P < 0.0001.

The bioluminescence signals of the PalmReNL-exosomes and -MVs were also analyzed in plasma samples collected at various time points (5 min, 2 h, 6 h, and 24 h) following retro-orbital (i.v.) or i.p. injections (Fig. 5h and Supplementary Fig. 5). The maximum bioluminescence signals were observed at 5 min in the i.v. injected reporter exosomes and MVs. In contrast, it took 2 h to reach the maximum levels of the bioluminescence signals in the i.p. injected reporter exosomes and MVs (Fig. 5h). The reporter MVs showed total area under the curve (AUC) for i.v. 3.2 × 10^8^ ± 5.7 × 10^7^ p/s·h; n=15; and for i.p. 3.2 × 10^8^ ± 2.5 × 10^7^ p/s·h; n=15; Fig. 5i). The reporter exosomes showed significantly higher bioluminescence signals compared to the reporter MVs (total AUC for i.v. 9.6 × 10^8^ ± 9.8 × 10^7^ p/s·h; n=15; *p*=0.0006; for i.p. 7.2 × 10^8^ ± 6.0 × 10^7^ p/s·h; n=15; *p*=0.0005; Fig. 5i).

To determine if the bioluminescence signals of reporter MVs could be enhanced by improved substrate availability *in vivo*^22^, in the next set of experiments we tested a Fz analog FFz. PalmReNL-MVs were i.p. injected as injection routes (i.v. vs i.p.) did not affect the AUC of PalmReNL-MVs (Fig. 5i). Because of its improved solubility, a higher dosage of FFz can be applicable, but here we injected the same dosage of 0.25 mg/kg as Fz to compare their sinsitivities. PalmReNL-MVs administered i.p. exhibited 6-fold more bioluminescence when FFz was i.v. injected as the substrate (2.5 × 10^6^ ± 3.4 × 10^5^ p/s; n=5; *p*=0.0005) as compared to Fz (4.2 × 10^5^ ± 1.1 × 10^5^ p/s; n=5; Supplementary Fig. 6).

### Bioluminescence signals in tumor-bearing mice decreased as the tumors grow

Despite their distinct sizes and cellular origins, the reporter exosomes and MVs derived from 4T1 cells behaved similarly *in vitro* and *in vivo* under the constraints of our experimental approach. However, the PalmReNL-MVs produced and retained significantly higher fluorescence signals compared to the PalmReNL-exosomes, in extracellular spaces as well as the intracellular environment after being taken up by cells likely due to their larger size. Tumor cell-derived MVs have been shown to play a key role in cancer progression by transferring oncogenic receptors to neighboring cells in the tumor microenvironment^30–32^. However, how tumor cell-derived MVs distribute throughout the body and contribute to metastasis formation has not been determined. In an effort to start deciphering the roles that MVs play under both physiological and pathological conditions, in the next sets of *in vivo* experiments we followed the behavior of reporter MVs in mice with or without mammary tumors.

Interestingly, the bioluminescence signals of PalmReNL-MVs were significantly lower in tumor-bearing mice compared to the healthy mice (Supplementary Fig. 7). Two weeks after tumor cell injection, the bioluminescence signal in PalmReNL-MVs injected i.p. was 2.2-fold lower [8.7 × 10^5^ ± 1.3 × 10^5^ p/s; n=4; *p*=0.002 (Supplementary Fig. 7b,d)], compared to control [1.9 × 10^6^ ± 1.6 × 10^5^ p/s; n=4 (Supplementary Fig. 7a,d)].

### Early induction of metastasis by multiple doses of tumor cell-derived MVs in mammary tumor-bearing mice

The decrease of the bioluminescence signal of the reporter MVs in the presence of tumors combined with the results of other *in vivo* and *in vitro* experiments suggest the involvement of EV-mediated signaling pathways in the modulation of mammary tumor progression, most probably at the level of the lungs. Therefore, in the next set of experiments we investigated whether or not MVs play any role in the induction of metastatic lesions.

One week after orthotopic injection of reporter 4T1-BGL cells constitutively expressing fLuc and eGFP, 90% of immunocompetent BALB/c mice developed detectable tumors in the mammary fat pad as revealed by bioluminescence imaging (BLI) (n=40). By two weeks after injection of the reporter 4T1 cells, the tumors were still growing steadily. Metastasis was evident in 50% of females that received multiple injections of MVs three weeks after the tumor formation, but only when the MVs were purified from 4T1 cells (Fig. 6i). The metastatic foci were detected *ex vivo* using D-luciferin as the substrate (Fig. 6j; arrow points to the fLuc bioluminescence signal detecting metastasis in the lung of a mouse treated with 4T1 cell-derived MVs).

**Fig. 6.**
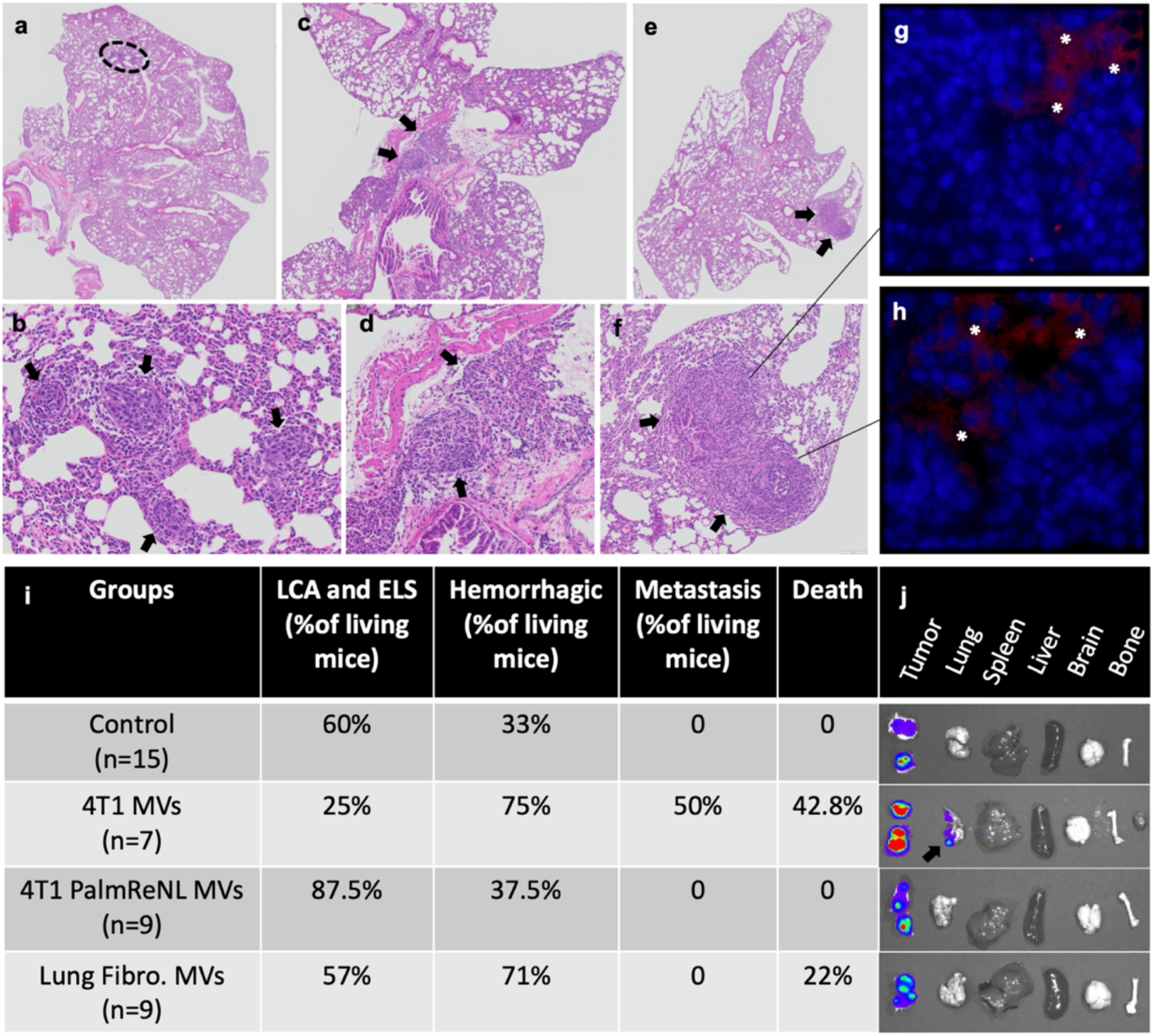
Promotion of early metastasis by multiple doses of tumor cell-derived MVs in mammary tumor-bearing mice. **a-f**, H&E of mouse lung tissues depicting the metastatic foci from two mice treated with 4T1 cell-derived MVs. The images in (a, c, e) are enlarged in (b, d, f). Black arrows indicate metastatic foci. **g, h** GFP-positive cells (stars) from the primary mammary tumors were observed only in areas near the metastatic foci. **i** The table depicts other findings including: hemorrhagic lungs, ectopic lymphoid structures (ELS) or lymphoid cell aggregates (LCA), and death. **j** In *ex vivo* BLI data, an arrow points to metastatic foci of 4T1-BGL cells detected in the lung using D-luciferin.

In addition, the metastatic foci were detected by histological analysis of lung sections following H&E staining (Fig. 6a-f). EGFP-positive cells were observed only in areas near the metastatic foci (Fig. 6g, h). Other histological findings included: hemorrhagic lungs were more apparent in mice treated with MVs purified from fibroblasts (71%) and 4T1 cells (75%) compared to control tumor-bearing mice; the appearance of ectopic lymphoid structures (ELS) or lymphoid cell aggregates (LCA)^33^ were predominantly present in lungs of mice treated with MVs isolated from 4T1 cells expressing PalmReNL (87.5%; Fig. 6i). No other significant pulmonary lesions other than intravascular evidence of systemic inflammation were observed.

The appearance of ELS and LCA structures within the lung suggested a strong inflammatory response due to the administration of PalmReNL-MVs. We confirmed this observation through performing immunohistochemistry of lung slides for the localization of F4/80 positive cells to detect foci of inflammation and particularly cells of the mononuclear phagocyte lineage^34^. As expected, cells positive for the F4/80 antigen were primarily associated with the lungs of mice treated with MVs derived from 4T1 cells expressing PalmReNL, localizing primarily to the bronchiolar epithelium and the pulmonary interstitium (Fig. 7e). Intriguingly, lungs treated with 4T1 cell-derived MVs demonstrated the most aggressive metastatic phenotype, while the F4/80 antigen did not show a remarkable immune response.

**Fig. 7.**
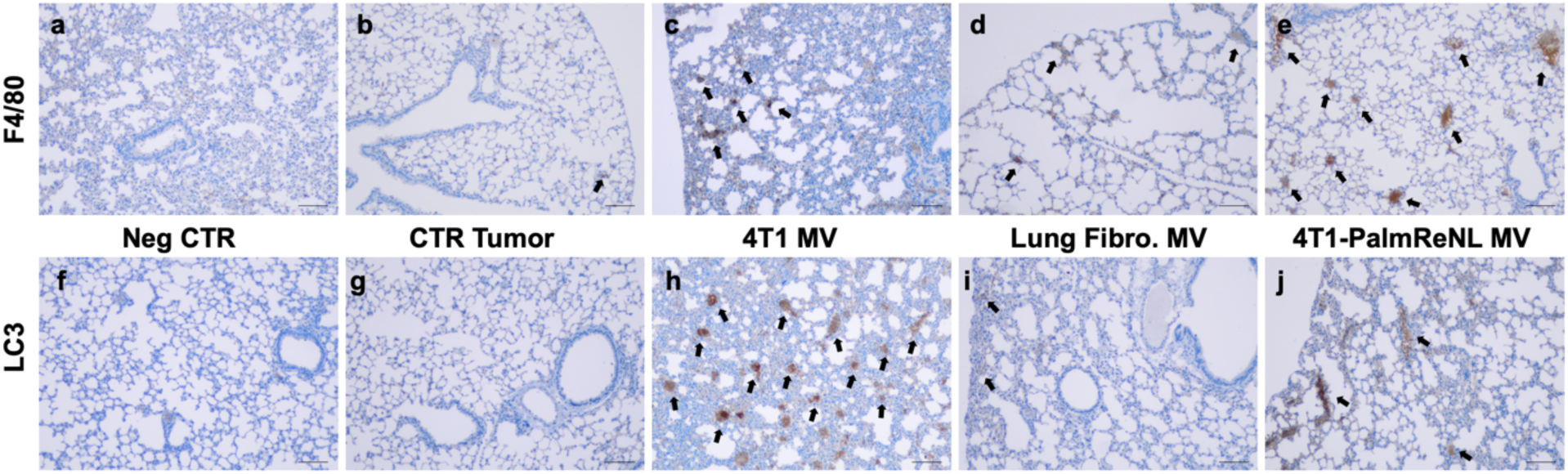
Immunohistochemistry detection of inflammatory foci and the LC3B protein expression. **a-e** Tissue sections of lungs from the different experimental groups stained with anti-F4/80 antibodies. **f-j** Tissue sections of lungs stained with anti-LC3B antibodies. Scale bars, 100 µm. Black arrows indicate F4/80 or LC3 positive regions.

This MV-mediated promotion of metastasis suggests that 4T1 cell-derived MVs may modify the tissue microenvironment within the lung in a way that potentiates survival of metastatic cells. Previous reports showed that EVs are released during stress and can ultimately metabolically reprogram adjacent cells, promoting survival, invasion, and metastasis^35,36^. We hypothesized that MVs may be potentiating the survival of cancer cells within the lung through a similar mechanism. To assess the metabolic health within the lung of MV-treated mice, we performed immunohistochemistry of the lung slides for the localization of MAP1LC3B (LC3), a key autophagy player that is often upregulated to promote cancer cell survival in the presence of metabolic stress^37^. Remarkably, the expression of the LC3 was upregulated in the lung tissue of mice that developed early metastasis following multiple injections of MVs purified from unmodified 4T1 cells (Fig. 7h).

### Autophagy knockout cell lines exhibited increased EV release

Because metabolic stress promotes EV release, we investigated the possibility that it may affect MV uptake and release. We tested this within U2OS cells, a well-characterized autophagy model cell line^38^, due to the stark LC3 upregulation seen in our lung metastasis models. Unfortunately, traditional tools used to induce metabolic stress, such as mitochondrial uncouplers or amino acid starvation, result in cell death within 24 hrs and we were unable to accumulate sufficient MVs within the conditioned media at this timepoint. Instead, we characterized the uptake and release of EVs within three different cell lines lacking autophagy-related genes (Atg2A/B, Atg5, and Atg9A; Supplementary Fig. 8), which are essential for the induction of LC3-dependent autophagy^39,40^. Cellular uptake of PalmReNL-EVs was characterized by assessing tdTomato fluorescence and measuring bioluminescence. Interestingly, authophagy KO cell lines had opposing effects on the uptake and release of EVs. Atg2A/B, Atg5, and Atg9A KO had reduced uptake of MVs derived from PalmReNL-4T1 cells at 24 h in U2OS cells as assessed by flow cytometry (Fig. 8d and Supplementary Fig. 9), while bioluminescence signals exhibited reliable results only at 2 h, possibly due to acid sensitivity of PalmReNL (Fig. 8g). The uptake of exosomes was reduced mainly in Atg2A/B and Atg5 KO cell lines (Fig. 8b). On the other hand, the release of MVs increased significantly in all the autophagy KO cell lines (Fig. 8j). The release of exosomes increased at the 72 h time point in the Atg2 KO cell line only (Fig. 8i).

**Fig. 8.**
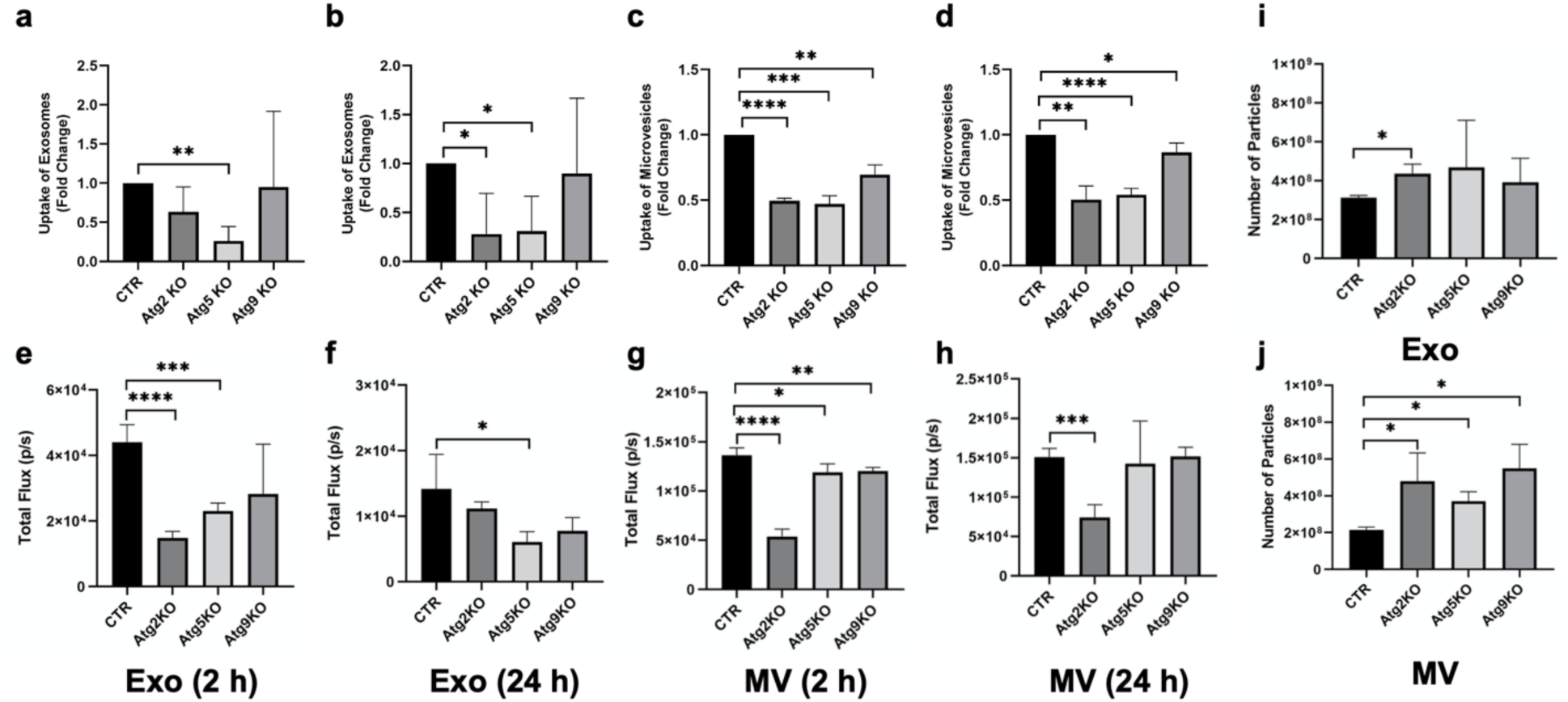
Decreased EV uptake and incresed EV release by blocking autophagy *in vitro*. Uptake of PalmReNL-exosomes or -MVs in control and autophagy knockout (KO) cell lines, analyzed by flow cytometry and measuring bioluminescence. **a-d** Flow cytometric analysis of cellular uptake of PalmReNL-EVs (tdTomato^+^) in U2OS Atg KO cells relative to control cells. The fold change of EV uptake was calculated using tdTomato fluorescence signals in KO cells compatred to the control (CTR). Error bars, SD (n = 3). **e-h** Uptake of PalmReNL-EVs determined by measuring bioluminescence signals. Error bars, SD (n = 4). **i** The release of exosomes assessed by NTA increased at the 72 h time point in the Atg2-KO cell line only. **j** The release of MVs increased significantly in all the autophagy KO cell lines. Error bars, SD (n = 3), *, P < 0.05; **, P < 0.01; ***, P < 0.001; ****, P < 0.0001.

## Discussion

Our BRET-based EV reporter presents a new tool for understanding the roles played by the distinct EV classes under physiological and pathological conditions. Using a novel PalmReNL EV probe, we visualized two major classes of EVs, exosomes and MVs, *in vitro* and *in vivo* under the constraints of current technologies for isolation and characterization of EVs. Our BRET-based EV reporter presents as a new tool for understanding the roles played by the distinct EV subtypes under physiological and pathological conditions. In addition to allowing us to determine with high sensitivity the *in vivo* biodistribution of exosomes and MVs, this EV reporter allowed us to decipher the possible roles of mammary tumor-derived MVs in mediating metastasis to the lungs. Although EV research remains restricted by current experimental limitations in separation of EV classes, the size and molecular markers of the EV subtypes in this study support the successful enrichment of exosomes and MVs in our experimental procedures^18,19^.

The addition of PalmReNL into the EV membrane did not appear to impair the biological functions of both exosomes and MVs, as demonstrated *in vitro* with the uptake experiments using various cell lines, and *in vivo* biodistribution studies in mice. Of note, PalmReNL-labeled MVs induced inflammatory responses in the lungs of the mammary tumor-bearing immunocompetent mice. Technical limitations with the detection of cellular uptake of PalmReNL-labeled exosomes gave the false impression of less uptake compared to MVs. However, we demonstrated the lack of sufficient sensitivity for the detection of fluorescence signals of PalmReNL in the recipient cells when tracking exosomes carrying CD63-mScarlet (Fig. 3c, f). Moreover, our analysis of single EVs by flow cytometry revealed that the fluorescence signals in PalmReNL-labeled exosomes were lower than those of PalmReNL-labeled MVs, possibly attributable to their size differences and/or symmetrical labeling of MV membranes. However, an equal number of exosomes showed higher bioluminescence signals than MVs. This result may indicate that PalmReNL can label more exosomes, while individual exosomes carry probe molecules not detectable by flow cytometry due to the detection limit. Therefore, analysis of bioluminescence signals may more appropriately reflect EV uptake in recipient cells.

We concluded that the Palm-based fluorescent EV probe is not ideal for tracking the fate of small EVs (i.e., exosomes) in recipient cells. Rapid diffusion of PalmReNL in endosomal network likely occurs after the cellular uptake of the reporter EVs, and the level of fluorescence signals goes below background. We demonstrated that the EV reporter may be processed through the endosomal-lysosomal pathway by using an acid-insensitive reporter. PalmGamillus-exosomes retained the fluorescence signals in the recipient cells compared to the PalmReNL-exosomes. Moreover, neutralizing the endosomal pH increased the bioluminescence signal intensity of PalmReNL-exosomes.

Recently, the heterogeneity of EVs was well-documented^41^. Therefore, it was not surprising that we found differences between the proliferation effects of exosomes and MVs. The signals of EVs differed most likely because of their various surface receptors and cargo. Additionally, recipient cells may have different activities of macropinocytosis and multiple modes of receptor-mediated endocytosis^42,43^. Interestingly, unlike other cell types tested, macrophage RAW 264.7 cells took up PalmReNL-MVs via clathrin- and caveolin-dependent pathways (Fig. 2m, q). In addition to phagocytic and macropinocytic EV uptake, this result corroborates findings that tumor cell-derived EVs are taken up efficiently by macrophages *in vivo*^44,45^.

In mice, the reporter-tagged exosomes and MVs showed similar *in vivo* biodistribution with respect to their blood circulation and organotropism, primarily towards the lungs. The analysis of the blood plasma also demonstrated that both types of EVs are quickly taken up and by 6 h after injection, their blood levels had decreased by half. Interestingly, the circulation time of the reporter MVs becomes shorter as tumors grow, possibly reflecting the involvement of immune responses^44,46^ as well as regulation orchestrated by the tumor and the establishment of premetastatic niches^45,47,48^. A study has described how MVs from metastatic melanoma cells enhance lung colonization of less aggressive, non-metastatic melanoma cells^49^. However, further work on MVs has mainly highlighted the ability of MVs to support primary tumor growth and survival^30,31,35^. The early appearance of metastatic foci observed in the present studies following multiple MV injections document potent far-reaching effects mediated by tumor cell-derived MVs and support their future potential as theranostic agents^11,50^. Our *in vitro* assays demonstrated that EVs can trigger biological responses in lung fibroblasts, AMSCs, macrophages, as well as in epithelial cells. Any one of these cells, if not all, could be involved in the preparation of the cell niche that favors metastasis to the lungs during repeated MV injections, a hypothesis that awaits confirmation. Importantly, in the present studies, MVs derived from normal tissues did not appear to contribute to the development of metastatic disease.

A better understanding of the EV-mediated systemic cross-talk between tumor cells and distant cells should aid in developing novel therapeutic approaches. For example, in lung and liver metastasis, exosomes exert their effect through immune cells and stromal cells^45,51^; in the bone, they mainly modulate local stromal cells, osteoclasts, and osteoblasts^52,53^. In liver metastasis of pancreatic cancer, macrophages play an essential role in receiving and relaying signals from tumor cell-derived exosomes^45^. Based on these studies, we hypothesized that MVs modulate metastatic behavior by orchestrating changes in the local tumor microenvironment as well as the systemic activation and recruitment of inflammatory cells at distant metastatic sites. However, results from our experiments demonstrated that the inflammatory cells, likely interstitial macrophages, appear to be more involved in prevention rather than in the promotion of early metastasis (Fig. 7). It is likely that the MV-mediated potentiation of metastasis is multifactorial. This is partially confirmed through the positiviely stained pockets of LC3 found within the lung tissue, which demonstrate difference in autophagic signaling within the lung. We hypothesize that tumor-derived MVs are modifying the tissue microenvironment within the lung through metabolic reprogramming. This alteration can promote proliferation, survival, and immune evasion of metastatic cancer cells in the regions they accumulate. However, further studies are required to characterize whether the upregulation of LC3 is a direct or indirect effect of MV administration. In addition, the influence of autophagy-related gene knockout on the release and uptake of EVs will need further characterization as well.

In summary, our new BRET EV reporter system enabled us to track EVs *in vitro* and *in vivo* with sufficient sensitivity. By combining non-invasive *in vivo* bioluminescence imaging with molecular and cellular analyses, we deciphered the possible role of MVs and LC3-associated mechanisms in early metastasis (summarized in Supplementary Fig. 10). Because of the complexity, EV-mediated signaling likely involves multiple pathways depending on variations of cell types and physiological/pathological conditions. Therefore, further extensive studies are needed to establish commonalities and functional differences of the EV classes. The ability to non-invasively image cancer-associated molecular markers will ultimately permit earlier detection and phenotyping of cancer, making possible the development of targeted therapies specific for individual patients.

## Methods

### Plasmid DNA constructs

All plasmids were constructed using standard PCR cloning protocols. The constructs were sequenced by GENEWIZ (South Plainfield, NJ) before using them for our experiments. For stable reporter gene expression, we constructed a Sleeping Beauty transposon^54^, in which the reporter genes were under control of the CAG promoter, by subcloning it into the multiple cloning site of the pKT2/CAGXSP vector^19^ through recombination cloning (In-Fusion HD Cloning Kit, Clontech). For the EV reporter, a palmitoylation sequence (MLCCMRRTKQ) of GAP-43^15,55^ was genetically fused to the NH_2_ terminus of NanoLuc^56^ (PalmNLuc), Red-eNanoLantern (ReNL^16^; Addgene plasmid #89536, gift from Takeharu Nagai), and Gamillus^26^ for EV membrane anchoring by PCR as reported previously. PalmNanoLuc and PalmReNL were amplified by PCR using (forward) 5’ -tggtggaattctgcagatagccgccaccATGCTGTGCTGTATGAGAAGAACCAAACAGGTCTTCACACTCGAAGATTTCGTTGGGGAC and (reverse) 5’ -cgccactgtgctggatTTACGCCAGAATGCGTTCGCAC, and (forward) 5’ -tggtggaattctgcagatagccgccaccATGCTGTGCTGTATGAGAAGAACCAAACAGGTGAGCAAGGGCGAGGAGGTC and (reverse) 5’ -cgccactgtgctggatTTACGCCAGAATGCGTTCGCAC, respectively. Human CD63 (Addgene plasmid #62964, gift from Paul Luzio) and mScarlet^25^ (Addgene plasmid #85042, gift from Dorus Gadella) were amplified using (forward) 5’ -tggaattctgcagatagccgccaccATGGCGGTGGAAGGAGGAATGAAATG and (reverse) 5’ -ccaccgctacctccacctcctagatctccCATCACCTCGTAGCCACTTCTGATACTCTTC, and (forward) 5’ -tggaggtagcggtggaggtggaagccaggatccgATGGTGAGCAAGGGCGAGGC and (reverse) 5’ -gccactgtgctggatTTACTTGTACAGCTCGTCCATGCCG, followed by combining these amplicons to generate CD63-mScarlet by overlap extension PCR. The sequence of Gamillus was synthesized as gBlocks (IDT) and a palmitoylation sequence was fused by PCR using (forward) 5’ -tggtggaattctgcagatagccgccaccATGCTGTGCTGTATGAGAAGAACCAAAC and (reverse) 5’ -cgccactgtgctggatTTACTTGTACAGCTCGTCCATGCCG.

### Cell culture

The labeling efficiency of the palmitoylated reporter was assessed in isolated fractions of exosomes and MVs from PalmReNL- or PalmNanoLuc-expressing 4T1 cells. The murine breast cancer 4T1 cells, murine macrophage Raw 264.7 cells, and primary mouse lung fibroblasts were cultured in DMEM supplemented with GlutaMAX (Gibco), 10% (vol/vol) FBS, and 1% penicillin/streptomycin. Mouse adipose-derived mesenchymal stromal cells (AMSCs) were cultured in α-DMEM supplementd with 15% FBS and 1% penicillin/streptomycin. Mouse primary lung fibroblasts and AMSCs were isolated as previously described^57^. Monoclonal human osteosarcoma cell (U2OS 3XFlag-HaloTag-Atg9A, see below) were cultured in RPMI media (Gibco/ThermoFisher, A4192301) supplemented with 10% FBS and 1% penicillin/streptomycin. All cell cultures were incubated at 37°C in a 5% CO_2_ atmosphere. Some cultures were treated with either methyl-β-cyclodextrin (MβCD; 10 mM), chlorpromazine (10 µg/mL)^58^, concanamycin-A (0.5 nM), or chloroquine (50 µM). All reagents were purchased from Sigma.

### Generation of the Atg knockout (KO) cell lines

A endogenously edited monoclonal human osteosarcoma cell line (U2OS 3XFlag-HaloTag-Atg9A) was used as the parental cell line (D.B. and J.S., manuscript in preparation) and subsequent knockout (KO) cell lines were generated by first ligating each genes corresponding sgRNA sequences (Atg2A: CGCTGCCCTTGTACAGATCG, Atg2B: ATGGACTCCGAAAACGGCCA, Atg5: AACTTGTTTCACGCTATATC, Atg9A: aggatatTCGAGAGAAGAAG, FlagTag: atggactacaaagaccatga) into the pX330-U6-Chimeric_BB-CBh-hSpCas9 backbone vector^59^ (Addgene plasmid # 42230; gift from Feng Zhang). These plasmids were co-transfected with GFP (pMAXGFP, Lonza), and single cells were sorted by FACS into 96-well plates, then screened and characterized by Western blot. In the case of the Atg9A-KO cell line, an additional flagTag sgRNA was added when the parental cell line was transfected.

### Cell proliferation assays

Cell lines were seeded in 96-well plates at a density of 5,000 cells per well, and 24 h later treated with 3.0 × 10^9^ exosomes or MVs. The viability of the cultures was determined 24 h and 48 h after EV treatments by using the CellTiter-Fluor cell viability assay kit from Promega (G6080) following manufacturers instructions. The filters in the fluorescence plate reader were set for AFC (380-400 nm excitation; and 505 nm emission).

### EV Isolation

EV-depleted FBS was prepared by 18-h ultracentrifugation at 100,000 g, 4^°^C^60^. 4T1, PalmReNL-4T1, or PalmNanoLuc-4T1 cells were seeded at 1.5 × 10^6^ cells per 100-mm cell culture dish. After 24 h the medium was replaced with EV-depleted medium and the cells were cultured for an additional 48-72 h. At the end of this culture period, MV- and exosome-enriched fractions were isolated as described previously^18,19^. Briefly, conditioned medium was centrifuged at 600 g for 5 min to remove cells and debris. The supernatant was centrifuged again at 2,000 g for 20 min at room temperature (RT) to remove apoptotic bodies. More dense MVs were separated from the less dense exosomes by centrifugation at 20,000 g for 60 min at 4^°^C, using a refrigerated microcentrifuge 5424 R (Eppendorf). Supernatants were filtered through 0.2 µm PES membrane filters (Nalgene, 725-2520) with pressure to remove large vesicles. Exosomes were collected by a size-based EV isolation method with modifications^19,61^ using 50-nm membrane filters (EMD Millipore, VMWP02500 or Whatman, WHA110603) with holders (EMD Millipore, SX0002500). Briefly, holders with 50-nm membrane filters were connected to a vacuum manifold (Qiagen), and washed with 5-10 mL of PBS buffer by applying vacuum. Next, the remaining exosome-enriched EV fraction in the supernatant were trapped on the membranes. When approximately 200-500 μL of sample remained, the concentrated exosome fraction was carefully collected. All EVs were aliquoted and stored at -80°C.

### Nanoparticle tracking analysis (NTA) and Zeta potential (ZP) measurement

EVs (exosomes and MVs) derived from 4T1 cells stably expressing PalmReNL, PalmNanoLuc, or without transfection were analyzed using the ZetaView Multiple Parameter Particle Tracking Analyzer (Particle Metrix) following the manufacturer’s instructions. EVs were diluted 100- to 1,000-fold with PBS or deionized water for the measurement of particle size and concentration. The ZP of EVs was measured after resuspending the EVs in deionized water to a concentration of 8.0 × 10^7^ particles/mL as previously reported^62^.

### Isolation of EV fractions by density gradient centrifugation

Conditioned medium from 4T1 PalmReNL cells was collected and centrifuged (600 g for 5 min, followed by another centrifugation at 2,000 g for 20 min). The supernatant was then loaded on top of a discontinuous iodixanol gradient (OptiPrep; Sigma D1556) of 5%, 10%, 20%, 30%, 40% and 50% layers^63^. The gradient was then ultracentrifuged at 100,000 g for 18 h at 4 °C. Six fractions were carefully collected from top to bottom, washed twice with PBS and pelleted by ultracentrifugation at 100,000 g for 1 h at 4°C. Efficiency, purity and concentration of the fractions were assessed by NTA and measuring bioluminescence signals.

### Flow cytometric analysis of surface phosphatidylserine (PS) in exosomes and MVs, and cellular uptake of EVs

For the analysis of EV surface PS, 100 µL of exosome- or MV-fractions derived from 4T1 cells expressing PalmReNL were stained with CellTrace Violet (CTV; Thermo Fisher, C34571), followed by removing free dye with a spin desalting column (BioVision, 6564). CTV-stained EVs were resuspended to a concentration of 3.5 × 10^9^ particles/mL in Hanks balanced salt solution (HBSS). 10x Annexin V binding buffer (0.1 M HEPES pH 7.4, 1.4 M NaCl, 25 mM CaCl_2_.) was added to the mixture (final concentration 1x), and incubated with Annexin V-APC (BioLegend; 8 ug/mL) to a final concentration of 0.7 µg/mL for 3 h at RT. At the end of the incubation period, the EVs were fixed with 2% paraformaldehyde (PFA, methanol-free, prepared with 1x Annexin V binding buffer; final volume 400 μL) for 20 min at RT. Immediately afterwards, the samples were run in the Cytek Aurora spectral cytometer in the MSU Flow Cytometry Core Facility. Signal threshold was set using the CTV signal to trigger off of detector V3 (violet), in order to discern EV signal from background. Buffer only and buffer with staining reagents were run as assay controls to reliably assess EV signals.

The uptake of PalmReNL-EVs by the Atg KO cell lines was also analyzed by flow cytometry. Cells were plated in 24-well plates at a density of 50,000 cells/well, 24 h later the cells were switched to EV-depleted medium, and 1 × 10^9^ EVs/well were added. After 24 h, medium was removed and the cells were washed with PBS, trypsinized and harvested. Immediately, the cells were fixed (4% PFA for 20 min at RT) and analyzed by flow cytometry for the fluorescence signal of tdTomato compared to unstained-cells (negative control) or 4T1 cells stably expressing PalmReNL (positive control).

Flow cytometry data were analyzed with FCS Express v7 (De Novo Software). Gates were drawn based on fluorescence minus one (FMO) controls.

### Fluorescence microscopy and Bioluminescence measurements

The uptake of PalmReNL EVs by murine macrophages (RAW 264.7), 4T1 cells, primary mouse lung fibroblasts, mouse AMSCs, or U2OS-Atg-KO cells was analyzed by fluorescence microscopy and measuring bioluminescence signals. The cells were plated in 96- or 24-well plates at a concentration of respectively 20,000 or 50,000 cells/well. 24 h later the cells were switched to EV-depleted medium and the reporter 4T1 cell-derived PalmReNL-exosomes or -MVs were added at a concentration of 3.5 × 10^9^ (for RAW 264.7, 4T1, lung fibroblasts, or AMSCs) or 1-3 × 10^9^ (for Atg-KO cell lines) EVs/well. At least three wells were analyzed for each treatment group (control, exosomes, MVs). The cultures were allowed to proceed for 24 h. At the end of the incubation period the uptake of the reporter was analyzed by fluorescence microscopy or measuring bioluminescence signals after adding furimazine (Fz; 25 μM) using a VICTOR Nivo Multimode Plate Reader or IVIS Lumina (PerkinElmer).

Phase contrast and fluorescence images of PalmReNL-4T1 cells, reporter EVs, or cells that were treated with the reporter EVs were taken using All-in-one Fluorescence Microscope BZ-X700 (KEYENCE) or DeltaVision Microscope (GE Healthcare Life Sciences). The cells were stained with 10 µg/mL Hoechst 33342 (H3570, Life Technologies) before microscopy was performed. All images were further analyzed using the ImageJ software (imagej.nih.gov).

### Western blotting

Whole cell lysates and equal numbers of EVs (3.75 × 10^8^ EVs) were derived from unmodified 4T1 and PalmReNL-4T1 cells, and mixed with 4x sample buffer (Bio-Rad) with β-mercaptoethanol (for detecting TSG101, ALIX, Flotillin-1, and anti-RFP) or without β-mercaptoethanol (for detecting CD63). Proteins were separated on a 4%–20% Mini-PROTEAN TGX gel (Bio-Rad) and transferred to a polyvinylidene difluoride membrane (Millipore, IPFL00010). After blocking with 5% ECL Blocking Agent (GE Healthcare, RPN2125) at RT for 1 h, membranes were probed with primary antibodies overnight at 4°C at dilutions recommended by the suppliers as follows: anti-Alix (Proteintech, 12422), TSG101 (Proteintech, 14497-1-AP), flottilin-1 (BD, 610820), anti-RFP (Rockland Immunochemicals, 600-401-379), CD63 (Thermo Fisher Scientific, 10628D, Ts63), Atg2A (Cell Signaling, 15011), Atg2B (Cell Signaling, 25155), Atg5 (Cell Signaling, 12994), Atg9 (Cell Signaling, 13509), followed by incubation with horseradish peroxidase HRP conjugated secondary antibodies at RT for 1 hour. The membranes were visualized with ECL select Western Blotting Detection Reagent (GE Healthcare, RPN2235) on ChemiDoc MP Imaging System (Bio-Rad).

### Dot blot analysis

Membrane orientation of PalmReNL in EVs was characterized as reported previously^15^. Both exosomes and MVs carrying PalmReNL (3 × 10^7^ EVs/µL) were 2-fold serially diluted. Aliquots were incubated in the presence or absence of 1% Triton X-100 for 30 min at 37 °C. After pre-wetting a polyvinylidene difluoride membrane (Millipore, IPFL00010) in methanol and equilibrating in transfer buffer, 2 µL of diluted EVs were dotted onto the membrane and blocked in 5% non-fat dry milk (RPI, M17200-1000.0) for 1 h at room temperature. The PalmReNL was detected using anti-RFP (Rockland Immunochemicals, 600-401-379) as described under Western blotting methods.

### Proteinase K protection assay

Exosomes and MVs carrying PalmReNL were split into 3 identical aliquots (3 × 10^7^ EVs/µL). Proteinase digestion was performed with 1 mg/mL proteinase K (Qiagen) in the presence or absence of 1% Triton X-100 for 30 min at 37 °C as reported previously^64^. At the end of a digestion period bioluminescence signals were measured after adding Fz (25 μM) using IVIS Lumina (PerkinElmer).

### Transmission electron microscopy

The samples were prepared as previously reported^61^, with slight modifications. Isolated EVs (PalmReNL-exosomes: 6 × 10^7^ EVs/µL, PalmReNL-MVs: 7 × 10^7^ EVs/µL) were fixed in 1% paraformaldehyde. A formvar-coated gold grid was kept in a saturated water environment for 24 h, and placed on a 50 µL aliquot of EV solution, and allowed to incubate for 20 min while covered. Next, samples were washed and blocked by placing each one face down on top of a 100 µL droplet of the following solutions: PBS (2x, 3 min), PBS / 50 mM Glycine (4x, 3 min), PBS / 5% BSA (1x, 10 min). A1:100 dilution of anti-RFP antibody (Rockland Immunochemicals, 600-401-379) in 5% BSA / PBS was used for labeling (1 h), followed by six washes in PBS / 0.5% BSA. Samples were incubated in a 1:50 dilution of donkey anti-rabbit immunogold conjugate (Jackson ImmunoResearch, 711-205-152) in 5% BSA / PBS (20 min) and washed in PBS (6x) and water (6x). The samples were negative stained with 1% uranyl acetate. Excess uranyl acetate was removed by contacting the grid edge with filter paper and the grid was air-dried. Samples were observed using a JEOL 1400 Flash Transmission Electron Microscope equipped with an integrated Matataki Flash sCMOS bottom-mounted camera. The 1400 Flash was operated at 100 kV.

### Bioluminescence pH titrations

Bioluminescence signals in exosomes and MVs (3 × 10^7^ EVs/µL) carrying PalmReNL were measured at room temperature (25°C) using a VICTOR Nivo Microplate Reader (PerkinElmer). EVs were incubated in the presence or absence of 1% Triton X-100 for 30 min at 37°C. The solutions consisted of 25 mM pH buffer, 125 mM KCl, 20 mM NaCl, 2 mM CaCl_2_, and 2 mM MgCl_2_ as reported^65^. The following buffers were used to adjust pH: pH 4.0 – 5.0: Acetate Buffer; pH 5.5 – 6.5: MES Buffer; pH 7.0 – 8.0: HEPES Buffer.

### *In vivo* tumor and metastasis studies

All procedures performed on animals were approved by the Institutional Animal Care and Use Committee of Michigan State University (East Lansing, MI). All mice were purchased from Charles River Laboratory. Eight week old female BALB/c mice with or without mammary tumors were used for the biodistribution studies of 4T1-PalmReNL-EVs (1.0 × 10^9^ particles/100 uL). For the induction of tumors, 2.5 × 10^4^ 4T1 cells constitutively expressing BSD-eGFP and fLuc (BGL) were orthotopically injected into the mammary fat pads of female mice under anesthesia. For the studies analyzing the development of metastasis, following tumor induction, MVs (3.0 × 10^9^ particles/100 uL) were injected into mice 3 times per week for 3 weeks (8 treatments in total). 4T1-BGL tumors were imaged (IVIS Spectrum system, see below) after i.p. injecting D-luciferin (3 mg/mouse in 100 μL PBS). Two weeks after tumor/metastasis induction the mice were imaged with the fluorofurimazine (FFz^22^) substrate (5 μg/mouse in 100 μL PBS), and the following day *in vivo* and *ex vivo* fLuc imaging were performed to analyze metastases. Immediately after, mice were sacrificed, dissected, and tissues were fixed (in neutral buffered formalin) and processed for histological analysis following paraffin embedding and H&E staining.

### Lung immunohistochemistry

Immunohistochemistry for detection of the LC3 protein and the macrophage marker F4/80 was carried out using standard protocols. Briefly, unstained sections of lungs or tumors were deparaffinized and rehydrated, and then incubated in the peroxidase blocking reagent (BioVision cat #K405-50). Antigen retrieval was perfomed by boiling the sections in sodium citrate for 20 min. To decrease background staining, the slides were incubated for 1 h in the mouse on mouse blocking reagent (Vector Labs MKB-2213-1), followed by overnight incubation with the primary antibodies (LC3B Cell Signaling 3868S rabbit polyclonal, 1:300; or F4/80 Cell Signaling 70076S rabbit monoclonal, 1:200). Next day, the slides were washed and incubated with One-Step HRP polymer (BioVision cat #K405-50) for 30 min at RT. The slides were then washed several times and then incubated with the DAB chromogen for 10 min at RT, followed by several washing steps and quick counterstain with Hematoxylin. Slides were then mounted and visualized under the upright microscope (Nikon).

### Bioluminescence imaging (BLI)

Bioluminescence analysis of the reporter exosomes and MVs was preceded by treatment of different concentrations of 4T1-PalmReNL-EVs (MVs and Exosomes) with 25 μM Fz or FFz (Promega). *In vitro* uptake or *in vivo* assays for the biodistribution of the reporter exosomes and MVs were imaged with IVIS Lumina or IVIS Spectrum systems (Xenogen product line of PerkinElmer). For *in vitro* assays, Fz was added to cultures of 4T1 cells, RAW 264.7 cells, lung fibroblasts, AMSCs, or U2OS-Atg-KO cells that were treated with the reporter exosomes or MVs prior to BLI. For *in vivo* imaging, mice were anesthetized with isoflurane using a SAS3 Anesthesia System (Summit Anesthesia Support) and an EVAC 4 waste gas evacuation system (Universal Vaporizer Support). Mice were injected retroorbitally (RO) or intraperitoneally (IP) with MVs or exosomes (1.0 × 10^9^ particles/100 µL) with either single or multiple injections (metastasis studies). Five min after intravenous EV injection, or 2 h after intraperitoneal (IP) EV injection, the mice were injected with either D-luciferin [150 mg/kg; IP], Fz (0.25 mg/kg; RO), or FFz (0.25 mg/kg; RO), and emitted photons were captured with IVIS as described previously^66^. Immediately after *in vivo* imaging, the mice were sacrificed and organs were excised. Organs were washed in PBS, treated with Fz, and imaged with the IVIS system. Bioluminescence signals were analyzed and quantified using the software program Living Image (PerkinElmer).

### Statistical analyses

All statistical analyses were performed with GraphPad Prism software (GraphPadSoftware). Screening results were analysed by one-way ANOVA followed by Tukey’s post-hoc test. Student *t* test was performed for all data set and *p* values were noted. Differences were considered to be statistically significant when the *p* value was less than 0.05.

## Author contributions

A.A.Z., G.I.P., M.K. conceived and designed the experiments. M.K. supervised the work. A.A.Z., G.I.P., D.B., B.D., A.M., V.T., L.K.T., M.P.B., A.W. executed the experimental work. A.A.Z., G.I.P., M.K. carried out the data interpretation and statistical analysis. J.H., J.R.W., T.A.K., M.H.B., J.S. provided reagents and technical advice. All authors contributed to the writing of the manuscript.

## Acknowledgements

We thank Mr. Nazar Filonov (Particle Metrix) for supporting our NTA and zeta potential measurement; Dr. A. Gilad for generously letting us use the C1000 Touch Thermal Cycler and ChemiDoc MP Imaging System; Dr. C. Mallett and Mr. J. Hix at the MSU IQ Advanced Molecular Imaging Facility; Ms. A. Porter at the MSU Investigative HistoPathology Laboratory; Dr. Loro L. Kujjo for critical reading of the manuscript. This work was funded by start-up funds from Michigan State University (MK. and JS). The schematic illustrations were created with Inkscape.

## Supporting Information

**Supplementary Figure 1.**
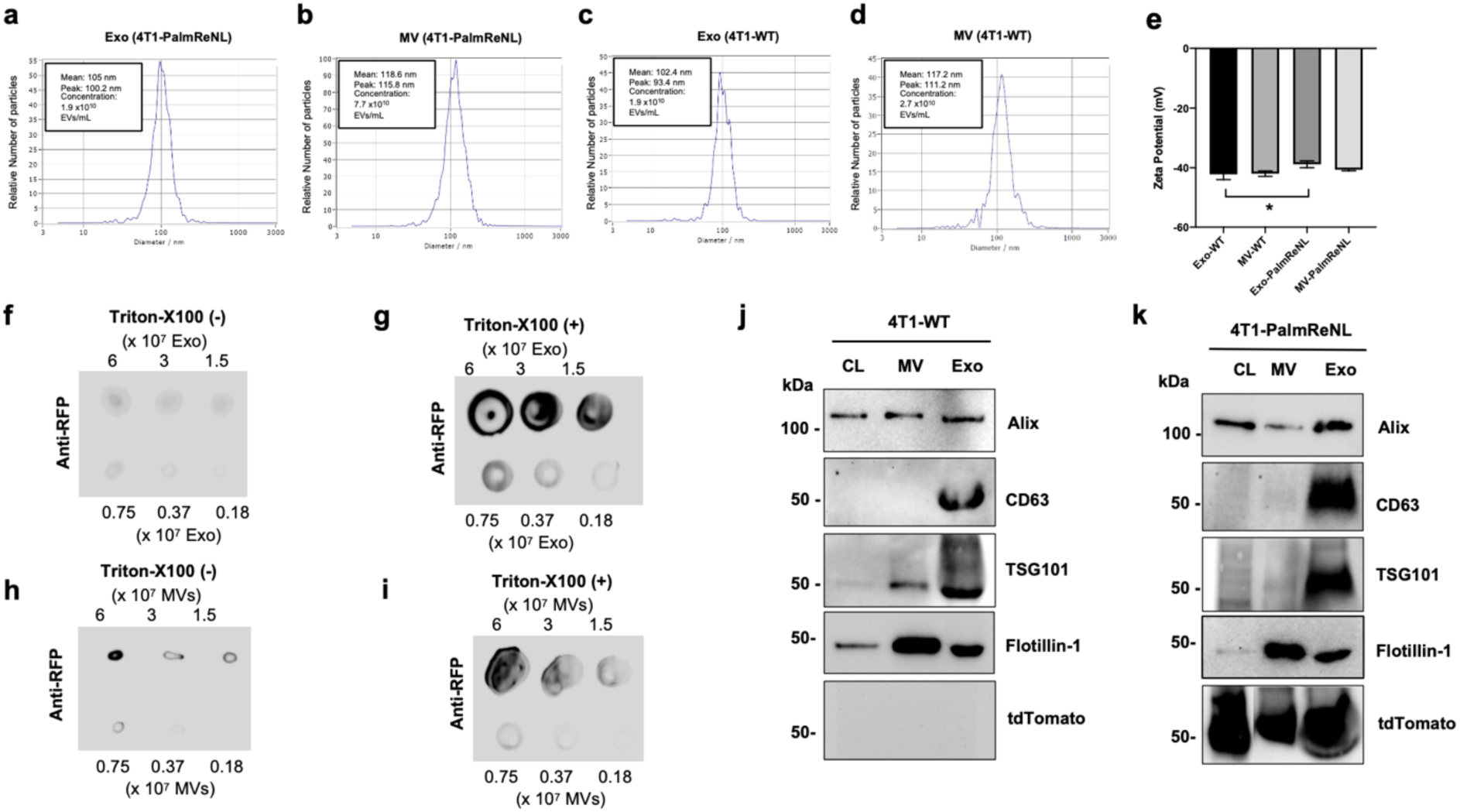
Characterization of 4T1 cell-derived PalmReNL-EVs. **a-d** 4T1 cell-derived PalmReNL-exosomes and -MVs were analyzed by nanoparticle tracking analysis (NTA). Exosomes and MVs derived from unmodified 4T1 cells were analyzed as control. **e** Determination of the Zeta potential revealed that the PalmReNL slightly shifted the surface charge of exosomes, but not MVs. Error bars, SD (n = 3), *, P < 0.05. **f, g, h, i** Dot blot assays for PalmReNL-exosomes and -MVs plus or minus Triton-X100. **j** Western blot analysis of exosome marker proteins of unmodified 4T1 cells, MVs, and exosomes. **k** Western blot analysis of exosome marker proteins of PalmReNL-4T1 cells, -MVs, and -exosomes.

**Supplementary Figure 2.**
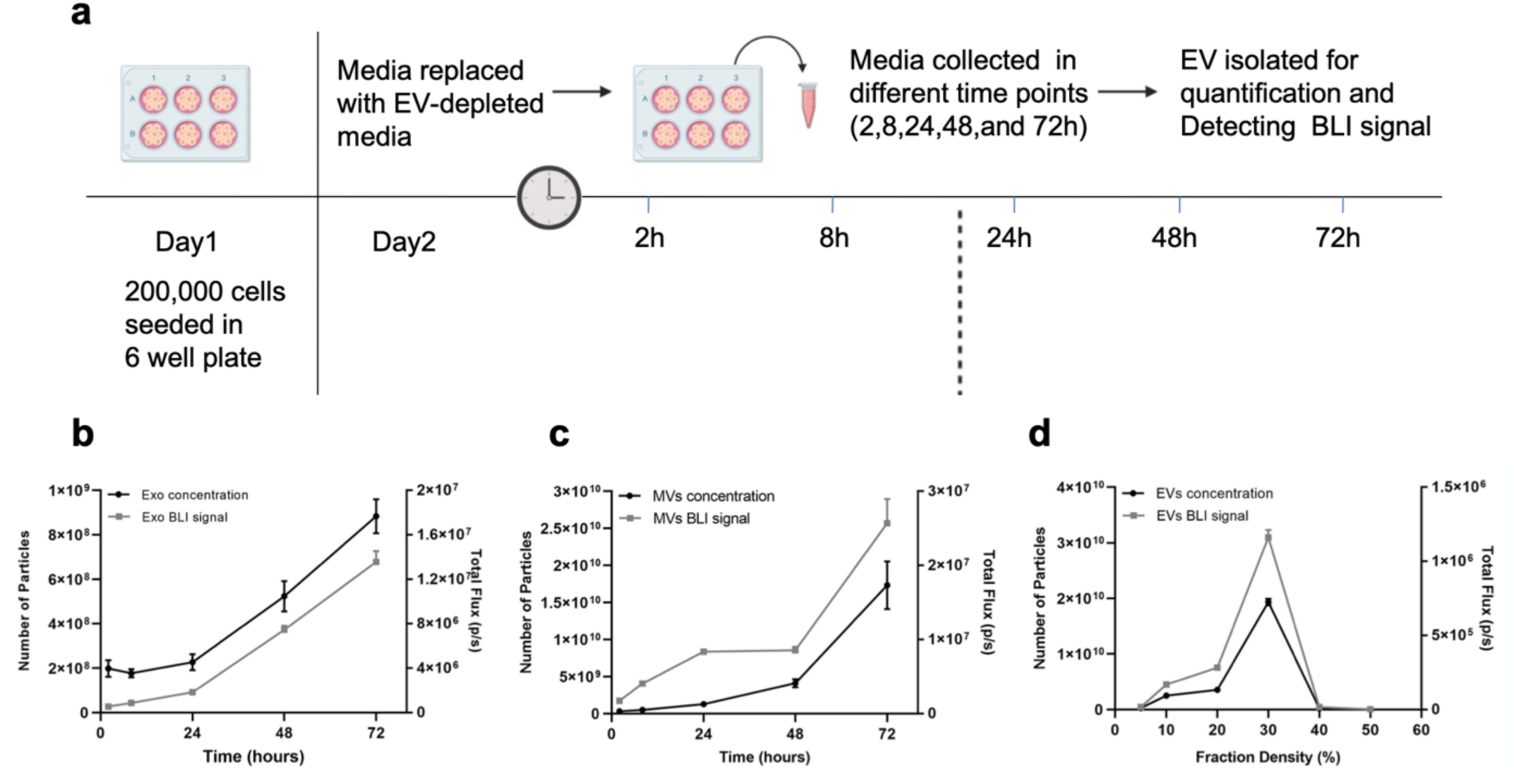
Characterization of EV release from 4T1 cells *in vitro*. **a** Schematic representation of the sampling protocol. **b, c** The bioluminescent signals correlated with the number of particles present. **d** The bioluminescent signals were also correlateed with the number of EVs purified from different EV fractions after density gradient. Error bars, SD (NTA, n=3; BLI, n=9).

**Supplementary Figure 3.**
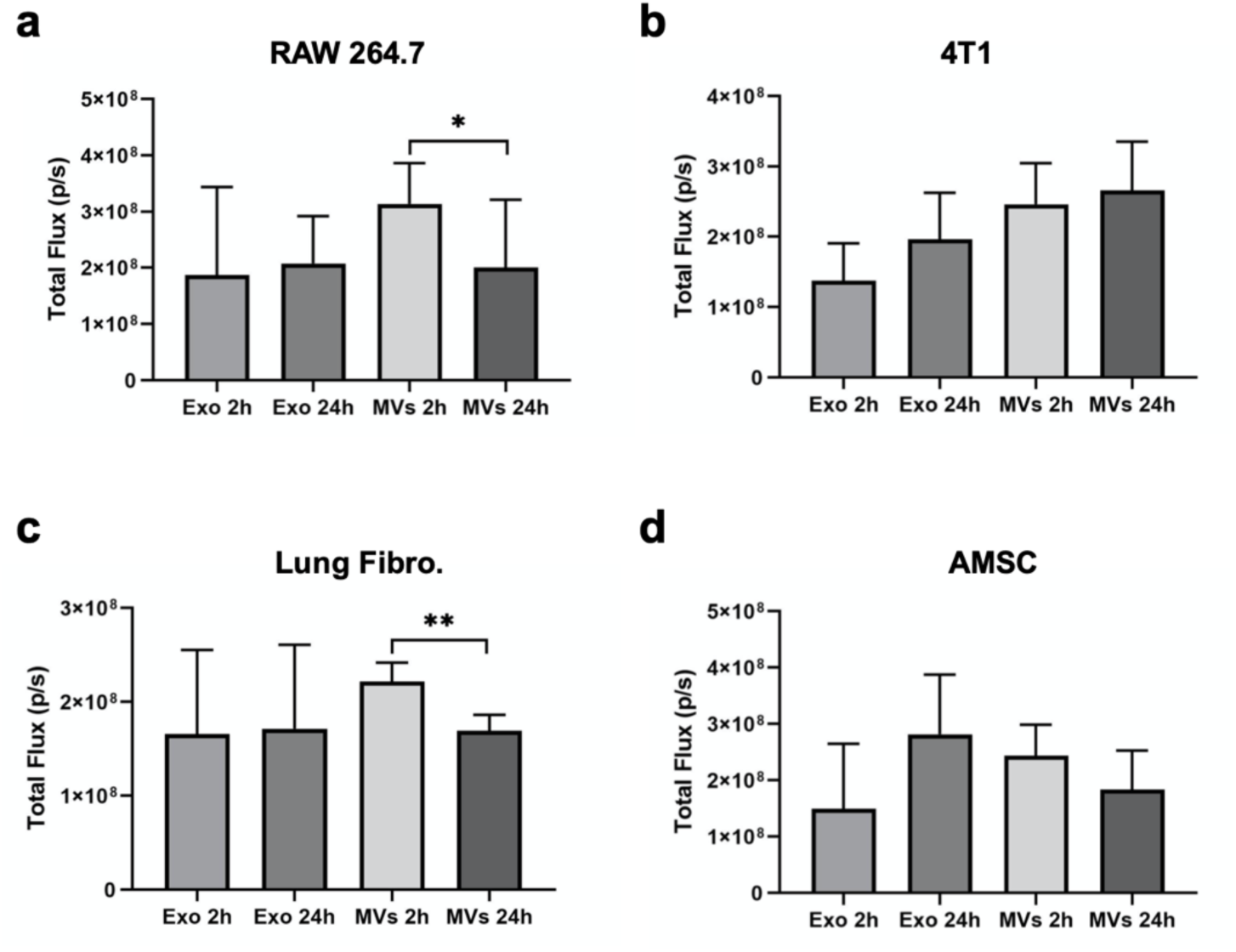
Short-term and long-term cellular uptake of 4T1 cell-derived PalmReNL-exosomes and -MVs assessed by measuring bioluminescent signals. **a** Macrophage RAW 264.7 cells. **b** 4T1 cells. **c** Mouse lung fibroblasts. **d** Mouse adipose-derived mesenchymal stromal cells (AMSCs). Error bars, SD (n = 8), *, P < 0.05; **, P < 0.01.

**Supplementary Figure 4.**
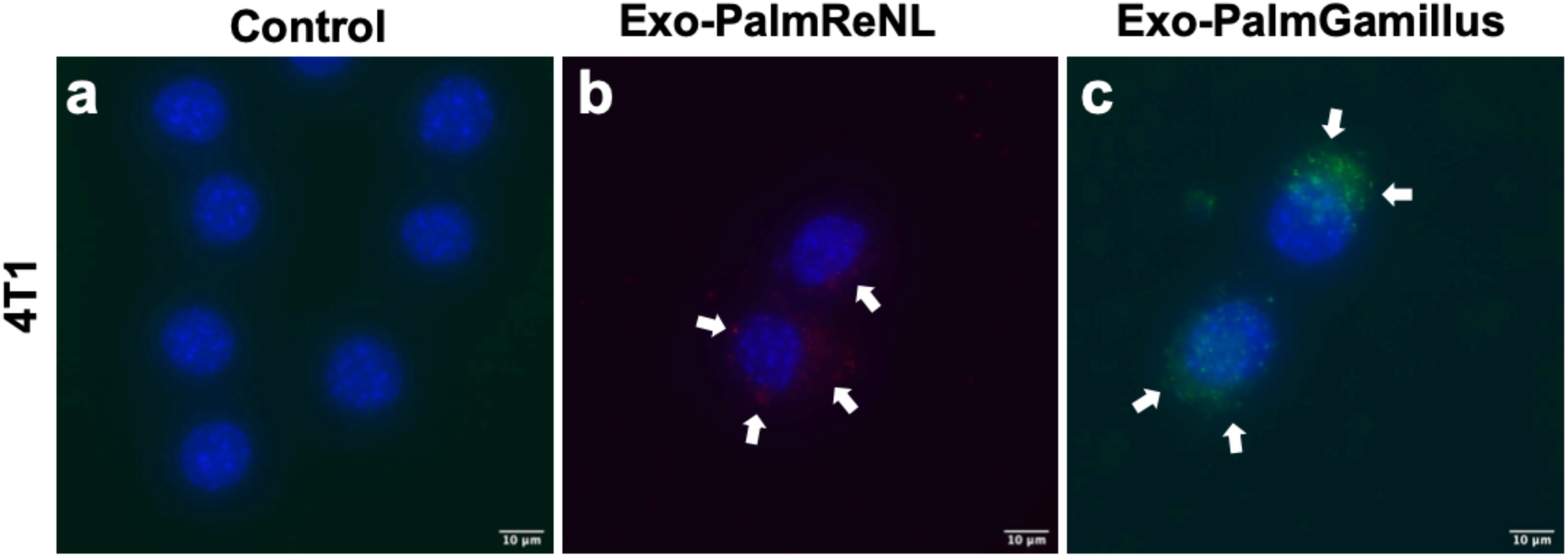
Retained fluorescence signals of PalmGamillus-exosomes in the recipient cells compared to the PalmReNL-exosome signal which was barely detected. Fluorescence microscopy images of 4T1 cells treated for 24 h with PalmReNL- or PalmGamillus-exosomes. White arrows, PalmReNL- or -PalmGamillus-exosomes.

**Supplementary Figure 5.**
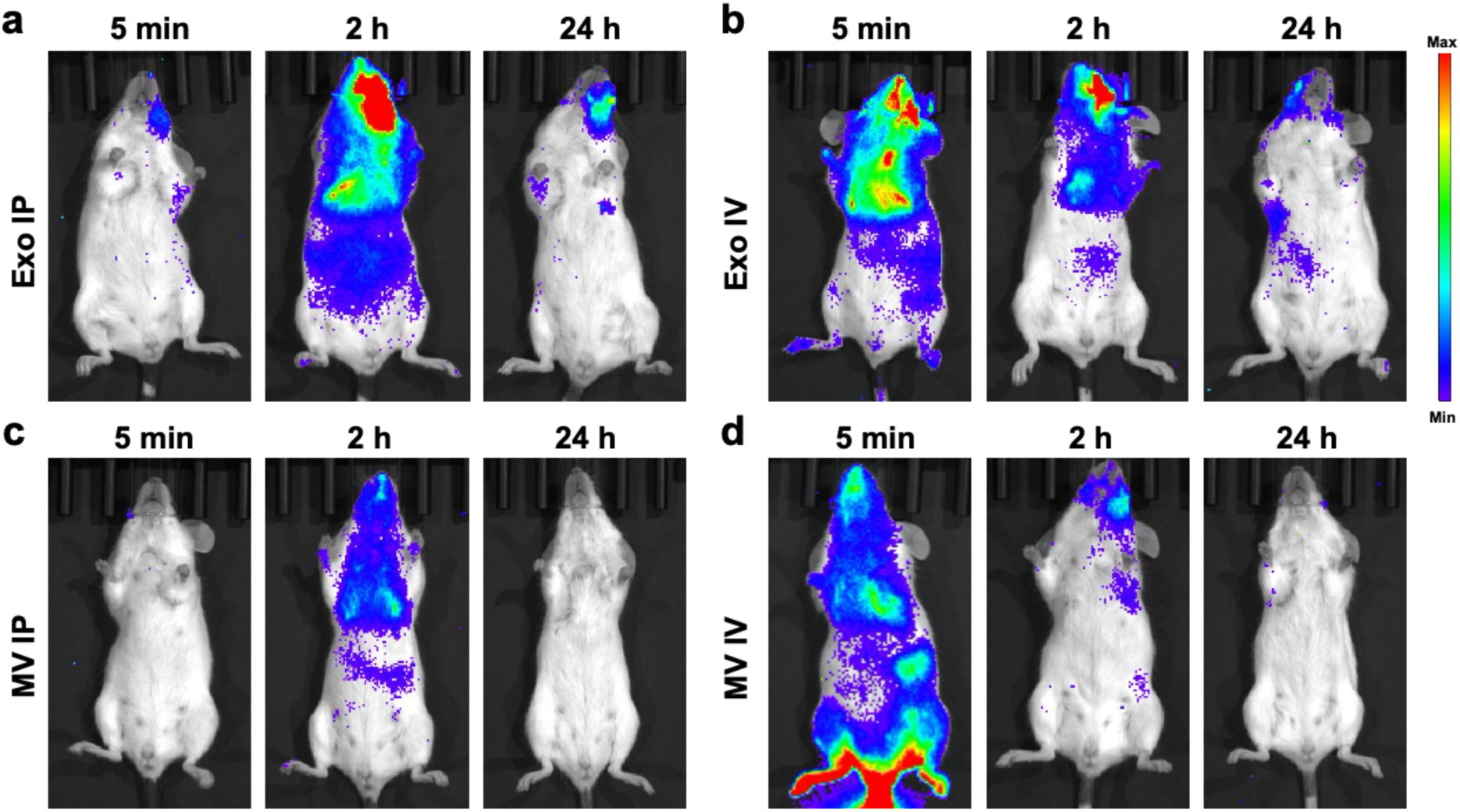
Dynamic biodistributions of PalmReNL-exosomes and - MVs following retro-orbital (i.v.) or intraperitoneal (i.p.) injections. **a-d** Representative biodistributions of the reporter PalmReNL-EVs (n = 3). *In vivo* BLI at 5 min, 2 h, and 24 h after injecting reporter PalmReNL-exosomes and -MVs. Furimazine (Fz) was administered retro-orbitally.

**Supplementary Figure 6.**
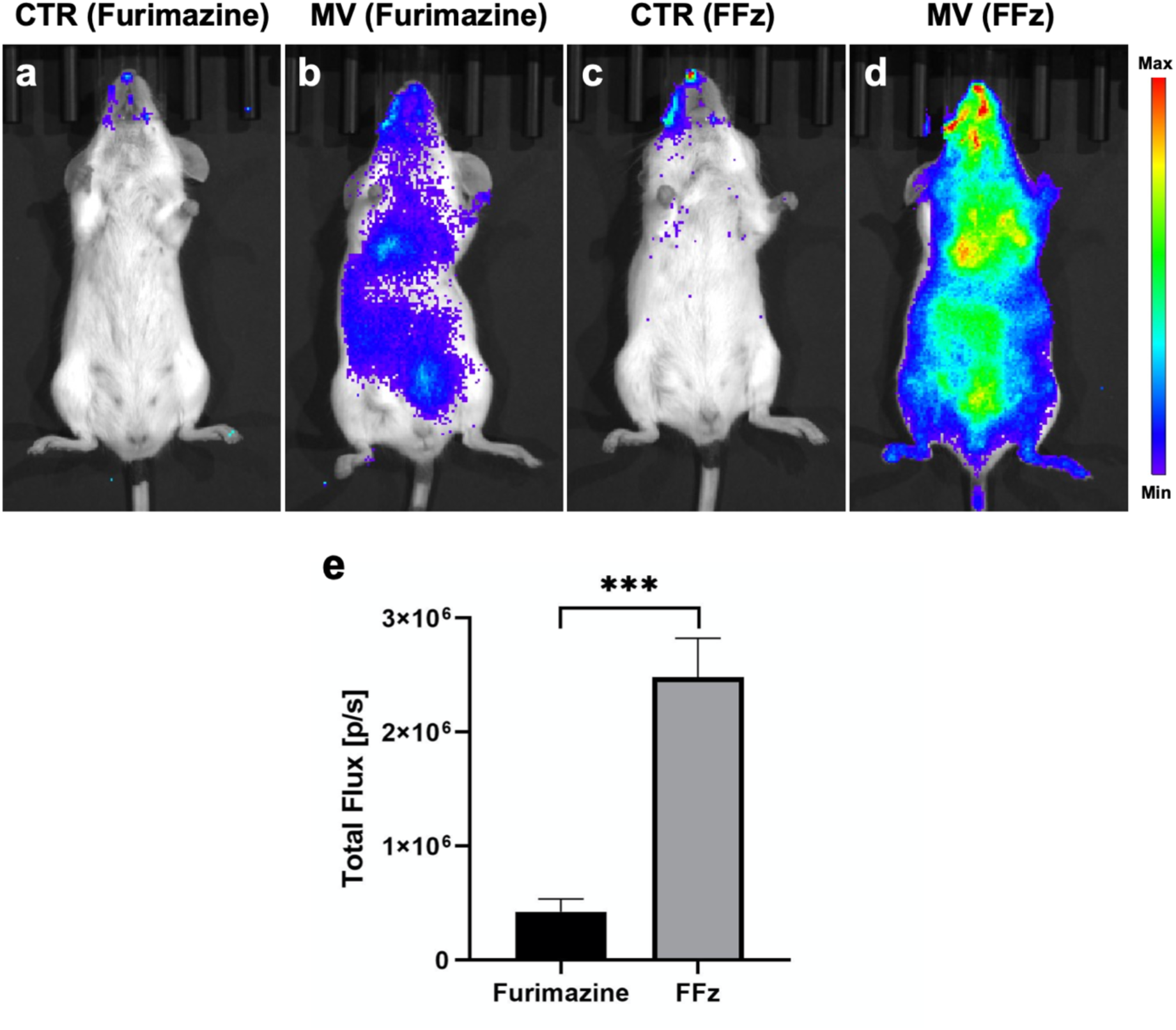
Enhanced bioluminescent signals of PalmReNL-MVs when Fluorofurimazine (FFz) was used as the substrate *in vivo*. **a, c** Control mice. **b, e** The bioluminescent signal of i.p. injected PalmReNL-MVs using furimazine as the substrate after. **d, e** The sensitivity of the reporter PalmReNL-MVs administered intraperitoneally was markedly increased when FFz was used as the substrate; Error bars, SEM (n = 5), ***, P < 0.001.

**Supplementary Figure 7.**
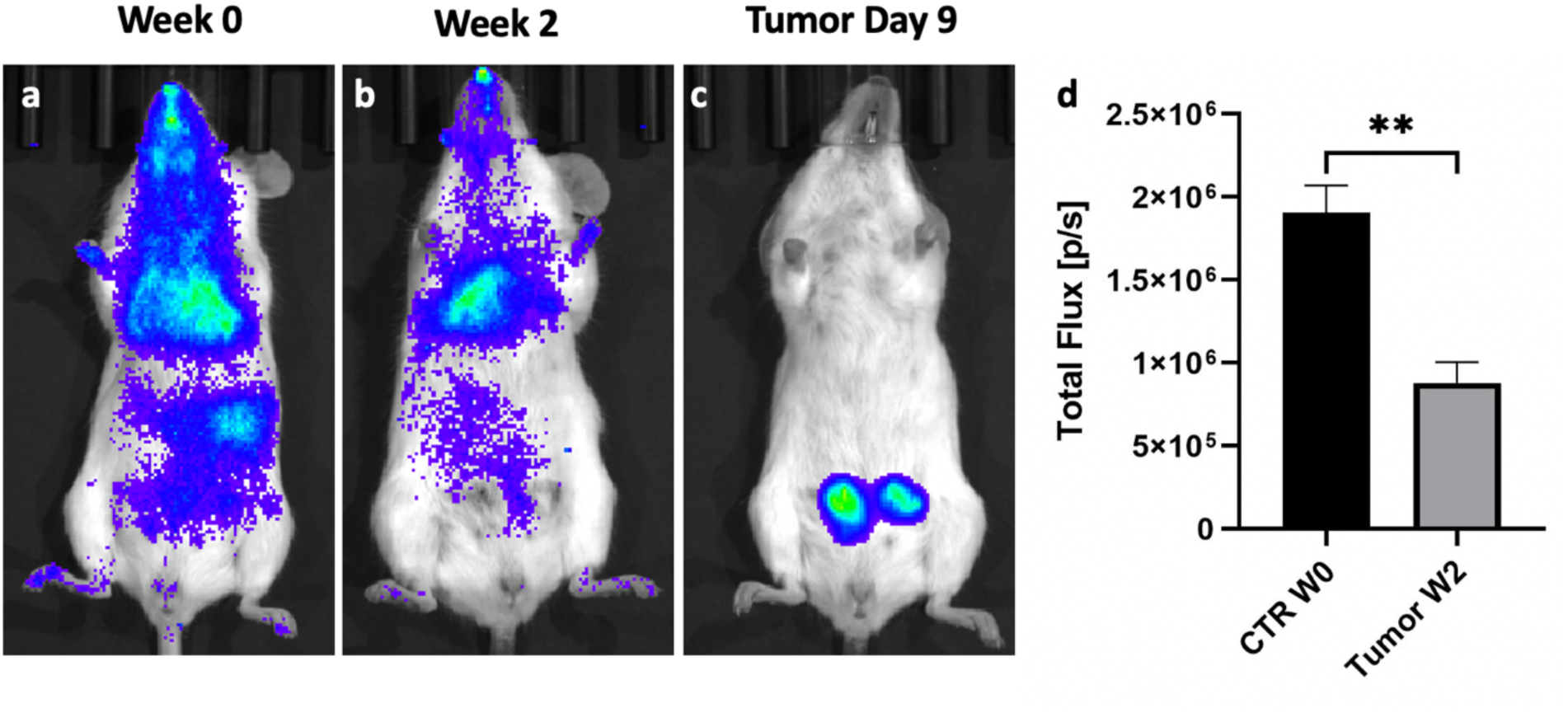
Lower bioluminescent signals of the reporter MVs-PalmReNL detected in tumor-bearing mice compared to the healthy mice. Bioluminescent signals of the PalmReNL-MVs in mice with and without tumors. **a** Control mice. **b** Mammary tumor-bearing mice two week after tumor implantation. **c** Mammary tumor growth was assessed by fLuc BLI. **f** Quantitative analysis of the bioluminescent signals of PalmReNL-MVs *in vivo* for each one of the experimental groups. Error bars, SEM (n = 4), **, P = 0.002.

**Supplementary Figure 8.**
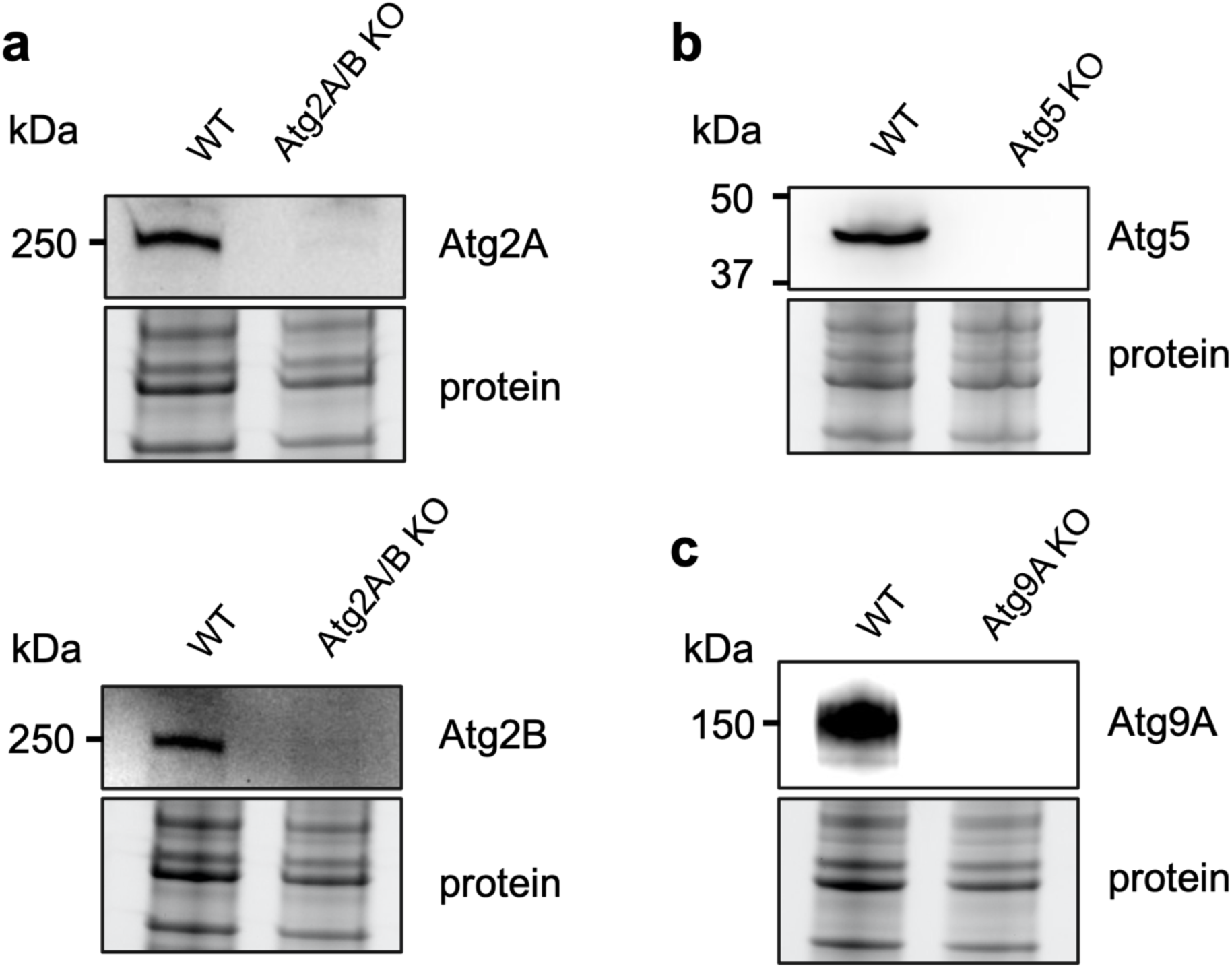
Characterization of Atg KO U2OS cells. **a-c** Western blot characterization of U2OS Atg KO cell lines. **a** Atg2A/B KO cells; **b** Atg5 KO cells; **c** Atg9A KO cells. Whole-cell lysates were shown as a loading control.

**Supplementary Figure 9.**
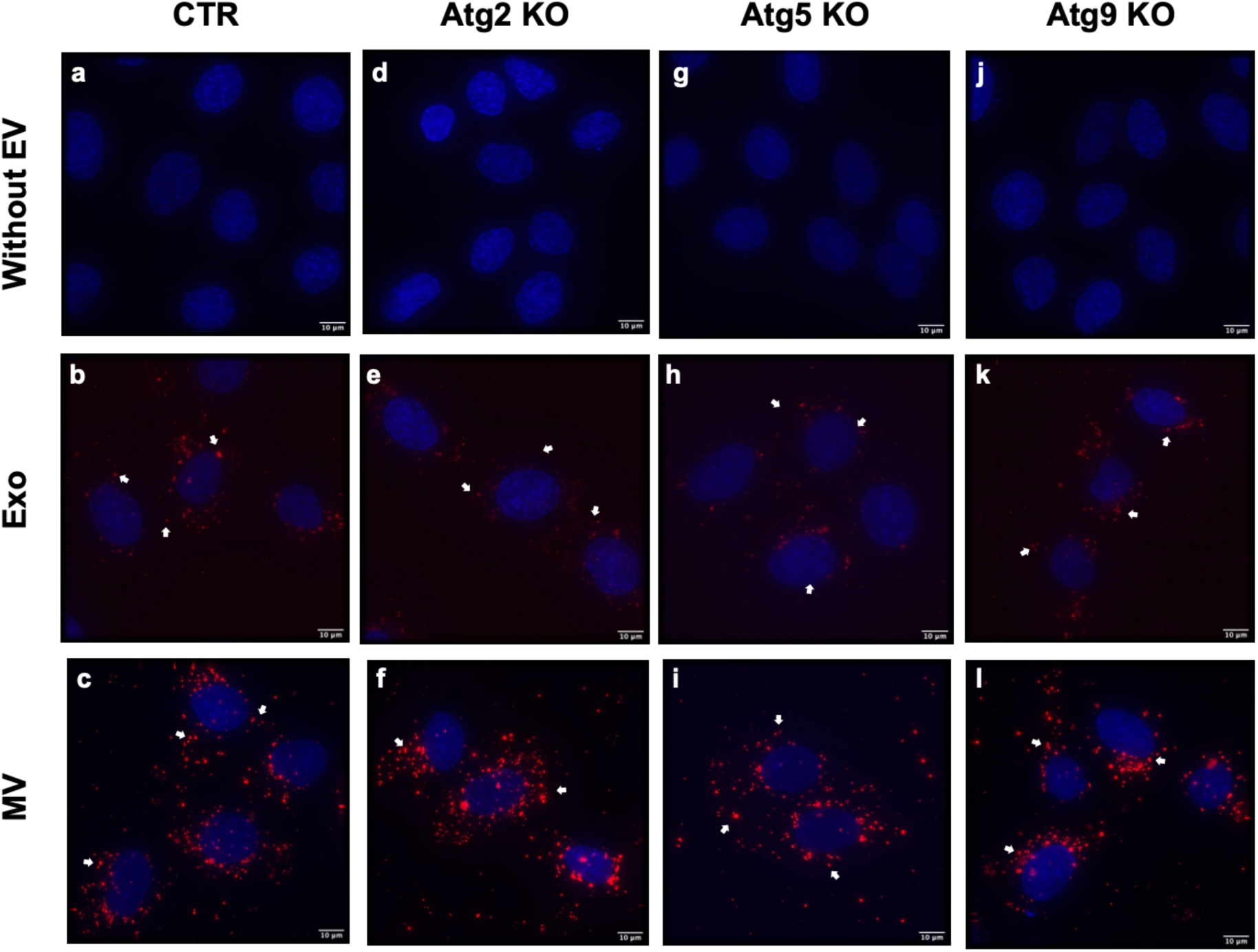
Autophagy involvement in cellular uptake of exosomes- and MVs-PalmReNL. **a-c** Control cells; **d-f** Atg2 KO cells; **g-i** Atg5 KO cells; **j-l** Atg9 KO cells. Punctate signals of RFP (red) were merged with nuclei stained with Hoechst 33342 (blue). Scale bar, 10 µm. Arrows indicate RFP signals in PalmReNL-exosomes and -MVs taken up by the recipient U2OS cells.

**Supplementary Figure 10.**
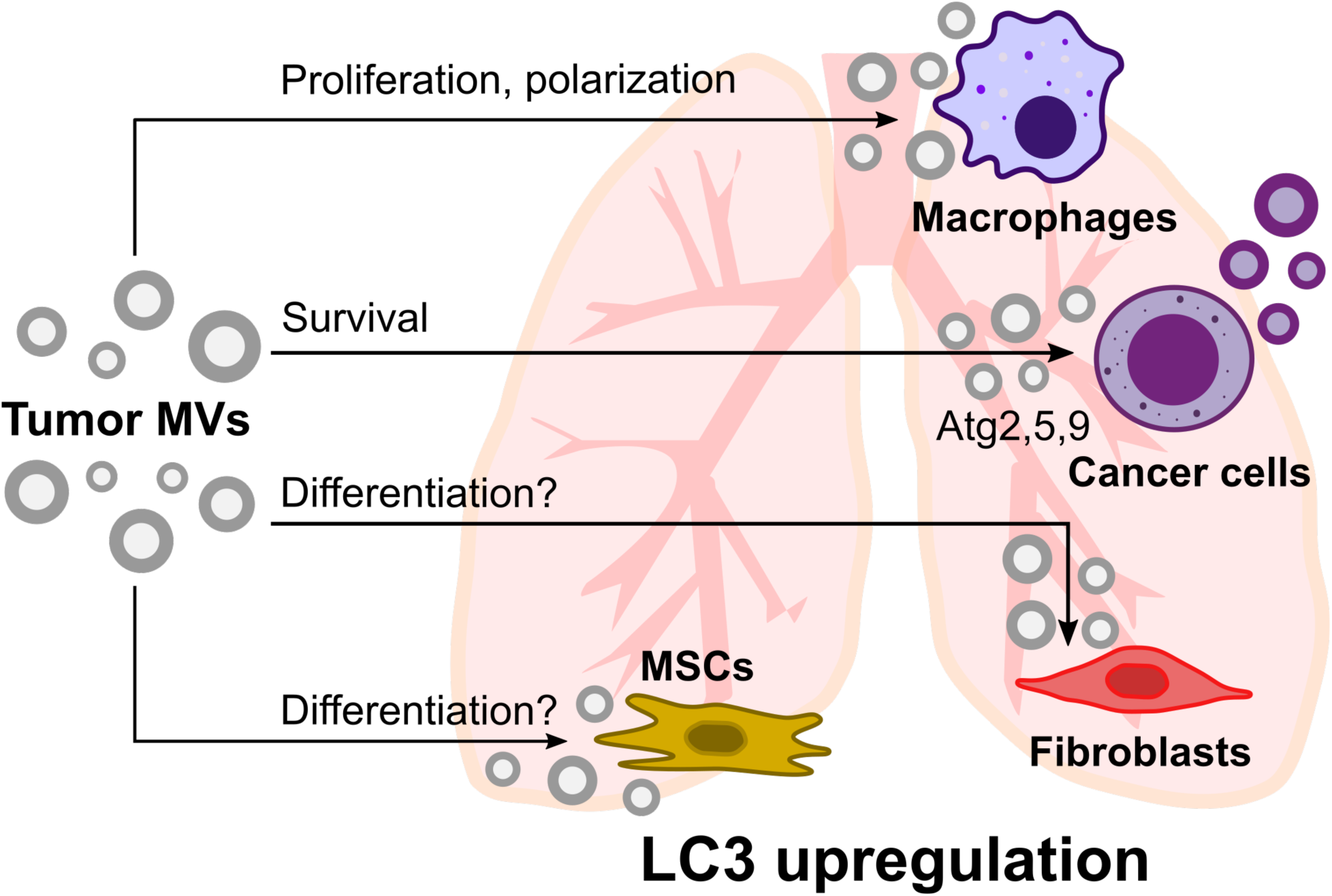
Schematic representation of metastatic cancer cell survival in the lung promoted by MVs from primary tumors. MVs released from primary tumors preferentially accumulate in the lung and affect various cell types, including macrophages, cancer cells, fibroblasts, and mesenchymal stromal cells (MSCs). Overall intercellular communication activates LC3 in the lung tissue and suppresses the anti-tumor response. Atg2, 5, and 9 proteins are directly and/or indirectly involved in metastatic cancer cells’ EV uptake and release.

